# Gene acquisition by giant transposons primes eukaryotes for rapid evolution via horizontal gene transfer

**DOI:** 10.1101/2023.11.22.568313

**Authors:** Andrew S Urquhart, Emile Gluck-Thaler, Aaron A. Vogan

## Abstract

Horizontal gene transfer (HGT) disseminates genetic information between species. The significance of HGT in eukaryotes is not well established, with evidence currently limited to isolated examples, typically absent of a mechanism. It has been proposed that mobile elements might be active agents of HGT in eukaryotes, as they are in prokaryotes. To test this hypothesis, we examined a gene cluster, which putatively contributes to formaldehyde resistance and is found within some members of the *Starship* superfamily of giant transposons. Remarkably, we found four instances where *ssf* has been independently acquired by distantly related *Starships*, and show that each acquisition event coincided with the element’s horizontal transfer (at least 9 HGT events in total). Our results demonstrate that acquisition of host-beneficial cargo by *Starships* primes fungi for rapid and repeated adaptation via HGT, revealing and elevating the role of HGT in eukaryotic biology.

## Introduction

Long thought to be a slow process taking place over millennia, adaptive evolution is now recognized as being able to operate over much shorter timescales [1]. Examples of rapid adaptation are now known from across the tree of life. In animals the textbook example of rapid adaptation comes from the peppered moth, which became darker in colour in response to increased pollution during the industrial revolution [2]. Other diverse examples include marine stickleback fish that have repeatedly adapted to colonise freshwater habitats [3], and house finches, which have evolved resistance to mycoplasma infection within the last 25 years [4]. Many organisms have rapidly evolved resistance to chemicals which humans have employed to control them, for example fungal pathogens developed widespread resistance to the azole fungicides used in agriculture and clinical settings occurred within several decades of there first use [5], bacteria have developed resistance to many antibiotics [6], the *Plasmodium* that cause malaria have developed resistance to antimalarials within 10 years of drugs deployment [7], and insects have become resistant to insecticides [8]. There are several mechanisms by which adaptation can occur rapidly. One is that natural selection can act upon existing standing variation in the population as in the case of stickleback fish [3, 9]. Another is that novel mutations can create major phenotypic change in a limited number of mutational steps. This is the case for fungicide resistance, which is due to simple mutations in the gene encoding the target of the fungicide [5, 10, 11]. Lastly, an organism may evolve a new phenotype by acquiring an existing genotype in a single step from another organism, known more simply as horizontal gene transfer (HGT) [12]. This last mode is well established as a driving force in the emergence of bacterial antibiotic resistance [13], but is poorly understood as a mechanism in eukaryotic evolution.

Despite lacking identified mechanisms enabling HGT in eukaryotes, there are a growing number of eukaryote-to-eukaryote HGTs implicated in adaptation. For example the whitefly *Bemisia tabaci*, obtained a gene from plants required to detoxify phenolic glucosides [14]. Similarly, aphids obtained the genes for carotenoid production from fungi [15]. In fish, an antifreeze protein gene horizontally transferred from herring to smelt [16, 17]. HGT seems to be particularly important in fungi. By the year 2000 there was growing, albeit inconclusive, evidence for fungal to fungal HGT, largely in the form of discordance between gene and species phylogenies [18]. More concrete evidence came by 2006 with the discovery that the gene encoding the plant host-specific toxin, *ToxA*, is 99.7% similarity between *P. tritici-repentis* and *S. nodorum* [19]. Such similarity was difficult to explain in the absence of HGT. Since then numerous additional examples have emerged including adaptation to cheese among fungi used in making cheese and numerous metabolic gene clusters [20–25]. However, unlike in prokaryotes the mechanisms underlying eukaryotic HGT are not yet well understood.

A better understanding of the genetic mechanisms underlying rapid evolution, including HGT, is of fundamental importance as it will help us predict if and when adaptive phenotypes might emerge. In fungi, this is a major problem as they threaten both human health and food security [26, 27]. *Starships* are a recently described superfamily of massive fungal transposons, which mobilise DNA within and between fungi [28, 29]. Some *Starships* carry genetic cargo (additional genes that do not encode functions for the transposition mechanism) that are adaptive for their fungal host. For example, the *Hephaestus* (*Hφ*) *Starship* of *Paecilomyces variotii* provides resistance to at least four toxic metal ions, zinc, cadmium, lead and arsenic [30]. The *Starships Horizon* and *Icarus* of the wheat pathogen *Pyrenophora tritici-repentis* contain the *ToxA* and *ToxB* genes, respectively, which increase pathogenicity on wheat [31]. Similarly, *Starship Voyager* is associated with a complex trade-off between saprotrophy and pathogenicity in the broad host range pathogen *Macrophomina phaseolina* [28]. Even the population-level frequencies of the *Starship Swordfish*, which is no longer presumed to be active, are associated with changes in climatic conditions in the wheat pathogen *Zymoseptoria* [32]. Additionally, many of these *Starships*, and/or their cargo, display signatures of HGT, implicating these massive mobile elements in the process of rapid adaptation. Furthermore, a much broader, yet uncharacterised, set of gene content is present in other predicted *Starships*, suggesting that a vast diversity of fungal phenotypes are mediated through HGT [28, 29].

Here, we test the hypothesis that *Starship* elements are a driver of rapid adaptation in fungi, via HGT, by investigating the evolution and function of the putative formaldehyde detoxification cluster of genes, *ssf*. We first found that *ssf* clusters have been acquired by four *Starships* belonging to distantly related families. Through a nucleotide-identity based approach, we show that each of these four elements containing the *ssf* cluster have horizontally transferred between species. We demonstrate that HGT of the *Starship Χρ* enabled rapid adaptation to formaldehyde in both *Aspergillus fumigatus* and *Paecilomyces variotii*, and confirm that this is controlled by the *ssf* cluster through gene knock-outs. Phylogenetic analyses confirm a single origin of the *ssf* cluster and its association to *Starships*, implicating inter-mobile element genetic exchanges as a catalyst for HGT. We also observe that a number of *ssf* clusters have integrated into genomic regions that are no longer associated with *Starship* elements, suggesting the birth-and-death processes of giant transposons shape the distributions of genes in fungal genomes beyond the lifespan of any single element. We argue that the *ssf* cluster is not unique, and that any genes which provide a similar selective advantage could be transferred by *Starships* and subsequently retained in a new host species. Thus, our results reveal that *Starship* dynamics drive the rapid emergence of adaptive fungal phenotypes, and elevate HGT from a phenomenon that rarely impacts eukaryotic genomes, to an impelling evolutionary force.

## Results

### The *ssf* region putatively responsible for formaldehyde resistance has been horizontally transferred by four different *Starship*s

We previously identified the *Starship Mι* from the Hephaestus-family as a *Starship* carrying a gene cluster encoding a number of proteins homologous to known formaldehyde detoxifying enzymes [29]. We name this region the *<u>S</u>tar<u>s</u>hip* formaldehyde cluster (*ssf*) (Figure 1A). The *ssf* region contains at least six and up to seven neighboring genes, the predicted functions of which are listed in Table 1. At least three of these were predicted to encode components of the glutathione-dependent formaldehyde detoxification pathway namely *ssfF*, *ssfA* and *ssfD* [34–37] (Figure 1B).

**Figure 1:**
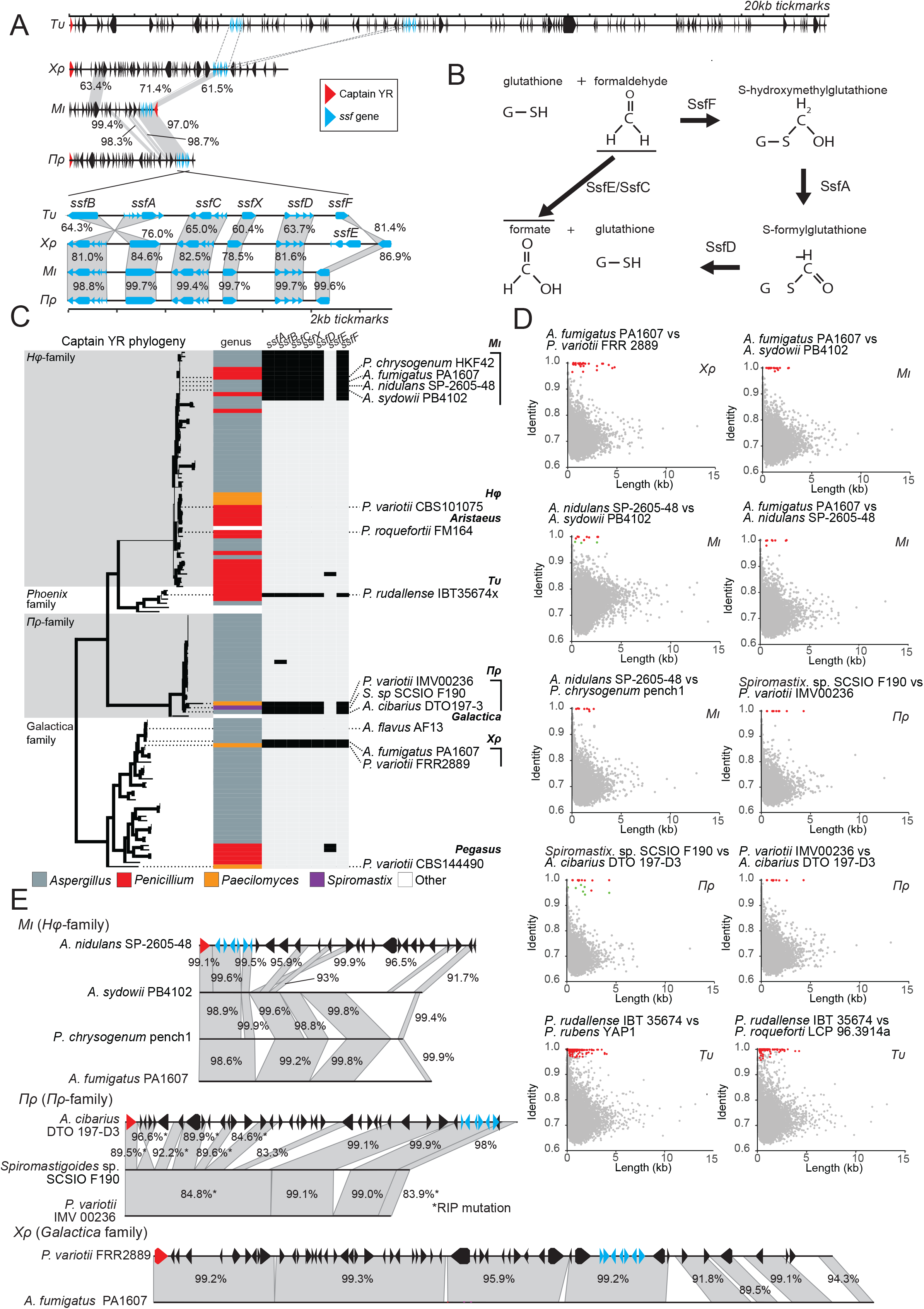
The *ssf* region putatively involved in formaldehyde detoxification has been horizontally transferred on four different *Starship* families. (A) A representative from each family of *ssf Starships* showing the position of the *ssf* gene region and below a comparison of the *ssf* region found in each *Starship*. The two copies of the *ssf* region in *Tυ* are nearly identical so only one of these is shown. Nucleotide identity is given in percentages based on Mauve alignments [55]. Dashed lines between *Tυ* and *Χρ* indicate homologous ssf regions too divergent to be aligned by Mauve. (B) Predicted roles of *ssf* genes in the glutathione-dependent formaldehyde detoxification pathway. (C) A phylogeny of *Starship* captain proteins from elements with the putative *ssf* cluster homologs and close relatives.(D) BLAST-all genome comparisons between genomes containing copies of the *ssf Starships*. Each point represents the best BLAST result from a single gene sequence. Points highlighted in red are genes within *Starships* we consider represent HGT based on high sequence identity in these genes compared to the remaining genes in the genome. Points highlighted in green as external to the *Starship* in one of the genomes. (E) Comparison of identified *ssf Starships* within each family reveals large amounts of highly similar DNA in each case. Nucleotide identity is given in percentages based on Mauve alignments [55]

**Table 1.**
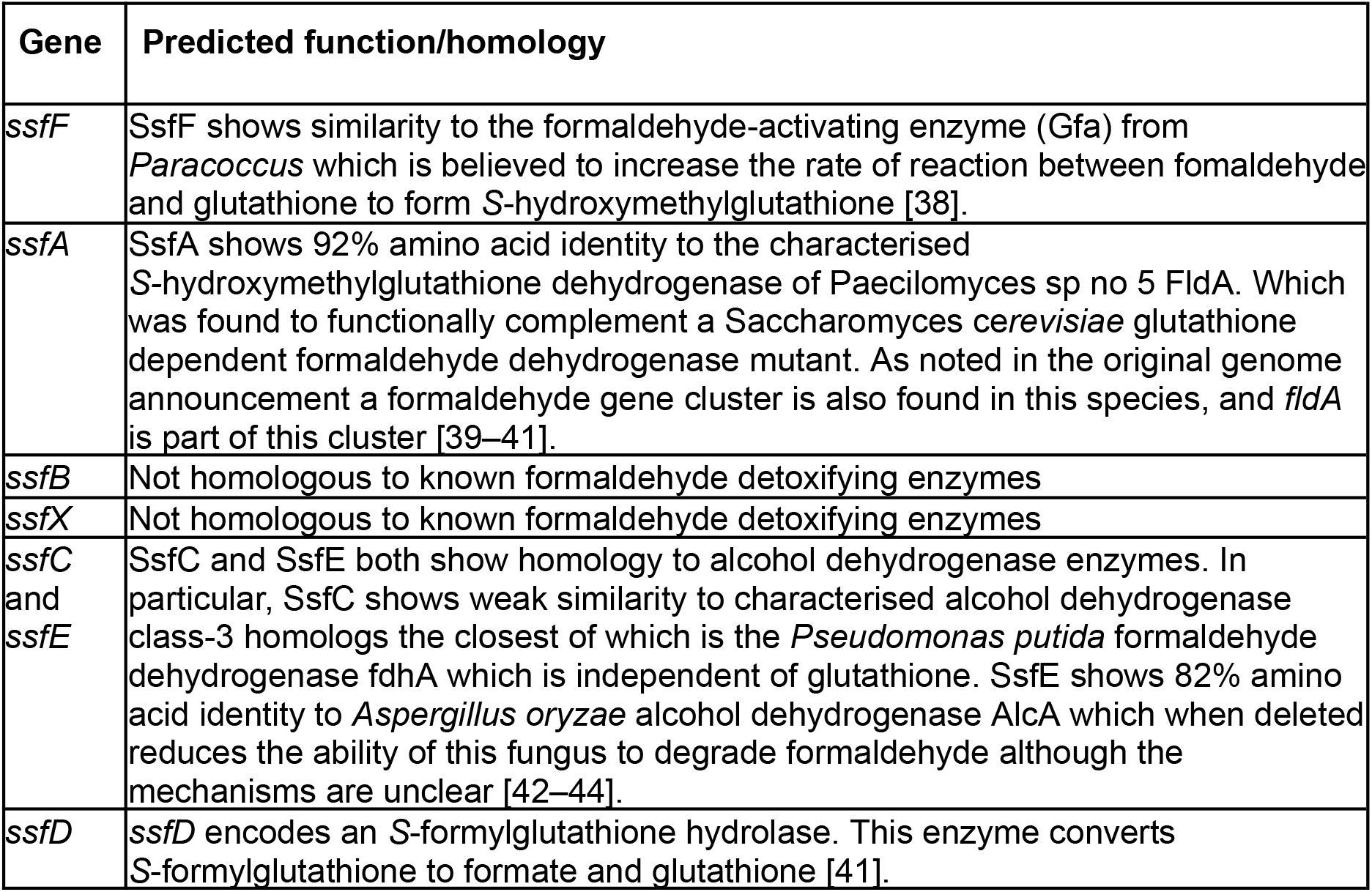

| Gene | Predicted function/homology |
| --- | --- |
| *ssfF* | SsfF shows similarity to the formaldehyde-activating enzyme (Gfa) from *Paracoccus* which is believed to increase the rate of reaction between formaldehyde and glutathione to form S-hydroxymethylglutathione [38]. |
| *ssfA* | SsfA shows 92% amino acid identity to the characterised S-hydroxymethylglutathione dehydrogenase of *Paecilomyces* sp no 5 FldA. Which was found to functionally complement a *Saccharomyces cerevisiae* glutathione dependent formaldehyde dehydrogenase mutant. As noted in the original genome announcement a formaldehyde gene cluster is also found in this species, and *fldA* is part of this cluster [39–41]. |
| *ssfB* | Not homologous to known formaldehyde detoxifying enzymes |
| *ssfX* | Not homologous to known formaldehyde detoxifying enzymes |
| *ssfC* and *ssfE* | SsfC and SsfE both show homology to alcohol dehydrogenase enzymes. In particular, SsfC shows weak similarity to characterised alcohol dehydrogenase class-3 homologs the closest of which is the *Pseudomonas putida* formaldehyde dehydrogenase *fdhA* which is independent of glutathione. SsfE shows 82% amino acid identity to *Aspergillus oryzae* alcohol dehydrogenase *AlcA* which when deleted reduces the ability of this fungus to degrade formaldehyde although the mechanisms are unclear [42–44]. |
| *ssfD* | *ssfD* encodes an S-formylglutathione hydrolase. This enzyme converts S-formylglutathione to formate and glutathione [41]. |

We have hypothesised that *Starships* disseminate traits that are beneficial for their genomic hosts because this might serve to increase their own chances of reproductive success [29]. We therefore examined the role of *Starships* in the horizontal movement of adaptive phenotypes using the *ssf* resistance cluster as a test case. We searched a large database of previously identified *Starships* for additional elements containing the *ssf* cluster [33]. This revealed that the *ssf* cluster is present in four distinct families of *Starship*: *Mι* (Hephaestus-family [30]), *Πρ* (Prometheus-family [33]), *Χρ* (Galactica-family [28]) and *Tυ* (Phoenix-family [28]) (Figure 1A).

Given that we previously showed that *Mι* likely horizontally transferred (not necessarily directly) between *Aspergillus* and *Penicillium* [29], we decided to investigate additional instances of horizontal transfer involving each of the families of *ssf*-carrying *Starships*. To do this we employed a “BLAST-all” approach that we have used previously to identify coding sequences with unusually high sequence similarity between two genomes [29, 30, 45] (Figure 1D). For *Mι*, in addition to being found within the genomes of *Aspergillus nidulans* SP-2605-48 [46] and *P. chrysogenum* “pench1”, as previously reported [30], *Mι* was also found within the genomes of several species in *Aspergillus* subsection *Versicolores* and *A. fumigatus* PA1607. From subsection *Versicolores* we selected a representative *Mι* from “*Aspergillus* sp.” PB4102 which we taxonomically identified as *A. sydowii* based on a phylogenetic comparison of the *RPB2* sequence to those examined by Jurjevic et al (Figure S1) [47]. “BLAST-all” comparisons between these four *Mι*-containing strains revealed a high level of sequence identity within *Mι* which is inconsistent with vertical descent from a common ancestor (Figure 1D and E). For example, BLAST of the *A. fumigatus* PA1607 gene regions vs the *A. sydowii* PB4102 assembly contigs returned hits for 7,652 genes with 20 genes returning a hit >100 bp and >96% identify (highlighted in red in Figure 1D). Of these 20 hits all were from genes located in the *A. fumigatus Mι* region (representing all of the *Mι* genes predicted). In contrast the remaining genes had BLAST hits with an average identity of 72.2%. Thus at least two HGT events have occurred between these pairs of *Aspergillus* species in addition to the HGT event reported previously, for a total of four events.

*Πρ* belongs to a recently described family of *Starships* which targets AT-rich sites [33]. We have identified three copies of *Πρ* containing the *ssf* cluster found within *Spiromastigoides* (previously *Spiromatix* [48]) sp. SCSIO F190 [49], and *P. variotii* IMV00236 [50], and *A. cibarius* DTO 197-D3 (Figure 1E). *Spiromastix* belongs to the order *Onygenales* and the latter two species to the order *Eurotiales*. Both the *P. variotii* and *Spiromastigoides* sp. *Πρ* are heavily affected by repeat induced point mutations (RIP), which is a fungal genome defence system that mutates repetitive DNA in the genome [51], and we thus consider these to represent immobile “derelict” *Starships*. Nevertheless, pairwise BLAST comparisons between these species showed highly conserved gene content within the *Spiromastigoides* sp. region in each species (Figure 1D). For example, BLAST searching the *Spiromastigoides*. sp. SCSIO F190 gene regions vs the *P. variotii* IMV00236 contigs returned a hit for 6176 genes with 10 genes returning a hit >100 bp and >100% identify (highlighted in red in Figure 1D). Of these 10 hits, all were located in the *Spiromastigoides Πρ* region (out of 18 *Πρ* genes predicted). In contrast, the hits in the rest of the genome showed only 72.3% average nucleotide identity. HGT of the *ssf* cluster must have occurred at least twice on *Πρ* including between the Eurotiomycete orders Onygenales and Eurotiales.

*Tυ* is a newly discovered *Starship* of the Phoenix-family [28] for which only one fully assembled element carrying the *ssf* cluster has been discovered. This is a ∼700 kb *Starship* in the genome of “*Pencillium sp.”* strain IBT 35674x which we identified as *A. rudallense* based on comparison of the RPB2 sequence to those examined by Houbraken et al. [52] (Figure S1). The massive size of the element, the largest *Starship* so far identified, means it is likely to be intact only in highly contiguous assemblies. However, at least two other *Penicillium* genomes contain a fragmented *Tυ*, *P. rubens* YAP1 and *P. roqueforti* LCP 96.3914a (genome published in [53]). BLAST comparison between *P. rudallense* and these genomes revealed signal for HGT both within and outside of the identified *Tυ Starship* (defined arbitrarily as genes with BLASTn matches with >100 bp length and >97% identity). The additional HGT highly conserved genes were found within a second *Tυ* element (which we designate *Tυ* haplotype 2) and on contig 8 where a degraded relict of *Tυ* is found (Figure 1D, Figure S2). Given the presence of a derelict starship we believe that the highly similar DNA on chromosome 8 represents DNA derived from this degrading *Starship*. Thus, HGT of the *ssf* cluster must have occurred at least twice on *Tυ* between *Penicillium* species.

Finally, in addition to being found within *P. variotii*, *Χρ* is found within certain *A. fumigatus* strains (including in PA1607, which also contains *Mι*) and this represents another HGT event (Figure 1D and E). BLAST searching *A. fumigatus* PA1607 gene regions as queries against the *P. variotii* FRR 2889 assembly contigs returned BLAST hits for 6,995 genes. Of these, 29 hits were over >100 bp in length and over 96% nucleotide identity (highlighted in red in Figure 1D). All of these hits were located within the *Χρ* region, in which 35 genes were identified in the copy found in PA1607. In contrast, the average nucleotide identity in the remaining BLAST hits was 72.5%. BLASTn searches also revealed the presence of *Χρ* in the genome of *A. flavus* strain B7001B, and while this assembly is highly fragmented, BLAST-all comparison between this genome and *A. fumigatus* PA1607 revealed strong HGT signal within *Χρ* (Figure S3).

In addition to looking at *Starships* carrying genes associated with formaldehyde tolerance, we also have identified an additional HGT of the original metal resistance *Hφ*-*Starship* via BLAST searches in the recently released genome of *Penicillium chermesinum* IBT 19713 [54] (Figure S4). This *Penicillium Starship* is nearly identical to the original *P. variotii Starship* except for the insertion of a retrotransposon (99.8% identical). The small amount of variation is mostly single nucleotide indels. Given that the *P. chermesinum* genome was generated using nanopore technology without additional polishing using Illumina reads, indel sequencing errors are expected and may account for the small amount of divergence observed [54].

### The transfer of a *Starship* carrying the *ssf* region underlies the evolution of formaldehyde resistance in both *Paecilomyces variotii* and *Aspergillus fumigatus*

We hypothesized that the observed horizontal transfer of multiple *Starships* containing the *ssf* region would enable the repeated evolution of formaldehyde resistance phenotypes in evolutionarily distinct lineages. To test this hypothesis, we focused on characterizing *Starship Χρ* because it contains all seven *ssf* genes and it is found in *A. fumigatus* and *P. variotii.* These two species are experimentally amenable with relatively large population-level sequencing datasets and represent both the putative donor (*Aspergillus*) and recipient (*Paecilomyces*) lineages of a *Starship-*mediated HGT event of the *ssf* cluster.

We first investigated a larger database of 512 publicly available short read archive datasets of *A. fumigatus* for the presence of *Χρ* [56, 57]. We found evidence for a full, non-truncated copy in just one strain, and were subsequently able to acquire this strain and 12 others that had no evidence of *Χρ* (Figure S5). Testing the formaldehyde tolerance of these *A. fumigatus* strains revealed that the strains lacking *Χρ* were unable to grow at 0.5 µl/ml formalin and showed either zero or restricted growth at 0.25 µl/ml formalin. In contrast, the *Χρ+* strain C-1-80s-1 grew relatively well at 0.25 µl/ml formalin and continued to show some growth at 0.5 µl/ml formalin (Figure 2A). We then assessed the distribution of *Χρ* within a set of six *Paecilomyces* strains available in our laboratory. These were five strains of *P. variotii* and one strain, FRR 5287, which belongs to its closest relative, *P. paravariotii* [45]. We found that *Χρ* is present only in *P. variotii* FRR 2889 and as a truncated copy in CBS 101075 (Figure S5). Testing the formaldehyde resistance of these strains revealed that the two *Paecilomyces* strains containing *Χρ* were more resistant than the four lacking *Χρ* (Figure 2B). The *Χρ*+ strains both showed growth on 6 µl/ml formalin whereas the *Χρ*-strains were all unable to grow on 2 µl/ml formalin. Together, these results demonstrate that the contribution of *Χρ* to formaldehyde resistance is maintained across HGT events.

**Figure 2:**
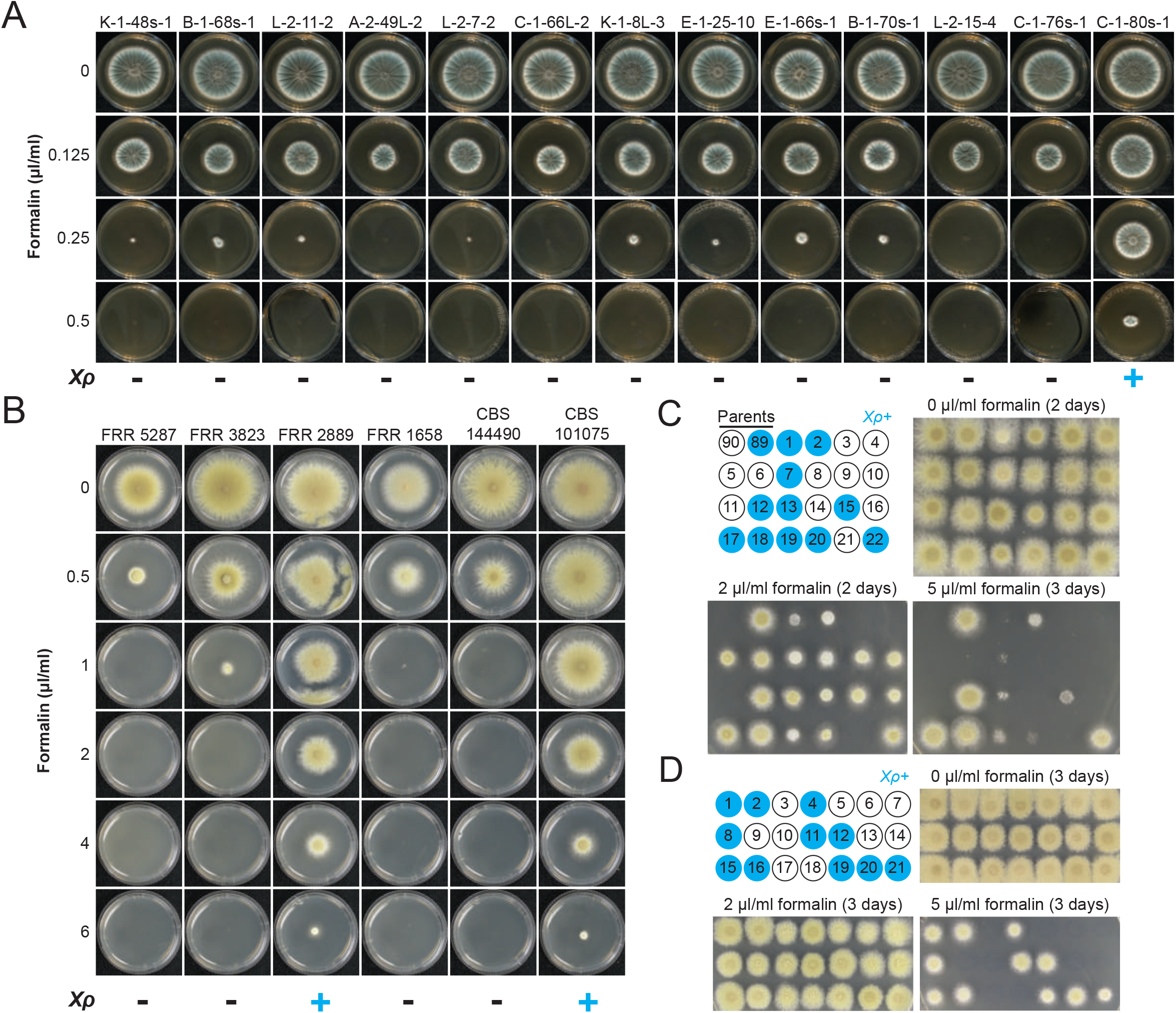
*Χρ* confers resistance to formaldehyde in *P. variotii* and *A. fumigatus*. (A) Wildtype *Aspergillus fumigatus* strains grown on varying formaldehyde concentrations showing increased formaldehyde resistance in the *Χρ*+ strain C-1-80s-1. (B) six *Paecilomyces* strains grown on varying formaldehyde concentrations. The two strains FRR 2889 and CBS 10175 containing *Χρ* are more resistant to formaldehyde than the four wildtype strains in which the region is absent. Plates were cultured at 37°C for 2 days. (C) Genetic segregation shows that *Χρ* contributes to increased formaldehyde tolerance in strains FRR 2889 and CBS 101075. Cosegregation was not apparent on 2 µl/ml formalin, whereas on 5 µl/ml formalin growth did cosegregate with *Χρ*. However, there was large variation with progeny #1 in particular germinating but showing poor growth which is barely visible. Those progeny that do not inherit *Χρ* show zero growth. (D) A back cross of progeny #6 from panel D to the *Xρ+* parental strain FRR 2889 produced progeny that all grew on 2 µl/ml formalin and on 5 µl/ml, segregated with the presence of *Χρ*. Plates were cultured at 30°C for 3 days.

To support the hypothesis based on this correlation that *Starship Χρ* is indeed responsible for the observed increase in formaldehyde resistance, we checked segregation patterns in *Paecilomyces* genetic crosses, by phenotyping progeny in the presence of formaldehyde and genotyping the progeny by PCR to assess the presence or absence of the formaldehyde region. Unexpectedly, segregation analysis using the progeny of a cross between FRR 2889 (*Χρ*+) and CBS 144490 (*Χρ*-) revealed that *Χρ* is only partially responsible for the increased formaldehyde resistance phenotype of these two isolates (Figure 2C). At 2 µl/ml formalin (a concentration at which the *Χρ-* isolates do not germinate) 18/22 progeny were able grow i.e. close to 75% instead of the expected 50%, which suggested that formaldehyde resistance might be linked to more than one independently segregating loci. The four progeny unable to germinate were all *Χρ-*. Increasing formalin to 5 µl/ml resulted in perfect segregation with the 11 *Χρ*+ progeny all germinating. However, the degree of growth varied widely with some *Χρ*+ progeny germinating but then showing very restricted growth (Figure 2C). Backcrossing progeny #6, which is *Χρ*-but presumably has the unmapped formaldehyde resistance loci given that it is able to grow at 2 µl/ml, to the *Χρ+* parental strain FRR 2889 results in progeny which all grow at 2 µl/ml and in which the presence of *Χρ* co-segregates with strong growth at 5 µl/ml formalin (Figure 2D). This indicated that at least one unmapped locus, together with *Χρ*, is contributing to increased formaldehyde resistance in these isolates.

### The *ssfB* gene makes the greatest contribution to formaldehyde tolerance

We next investigated the genetic basis of *ssf*-mediated formaldehyde resistance and found that genes within this cluster contribute unequally to this phenotype. First, the coding region of each individual *ssf* gene was replaced with that of GFP such that GFP expression came under the control of each native *ssf* promoter, thus allowing induction in response to formaldehyde to be visualised. Some induction was present in *ssfA*, *ssfB*, *ssfC*, *ssfD* and *ssfF* after the strains were germinated overnight in liquid media containing 0.1 µl/mL formalin with the greatest induction observed in *ssfA*, *ssfB* and *ssfF* (Figure 3A, S6). The response of native *ssf* promoters to formaldehyde supports a role for this region in in formaldehyde concentrations in the environment

**Figure 3:**
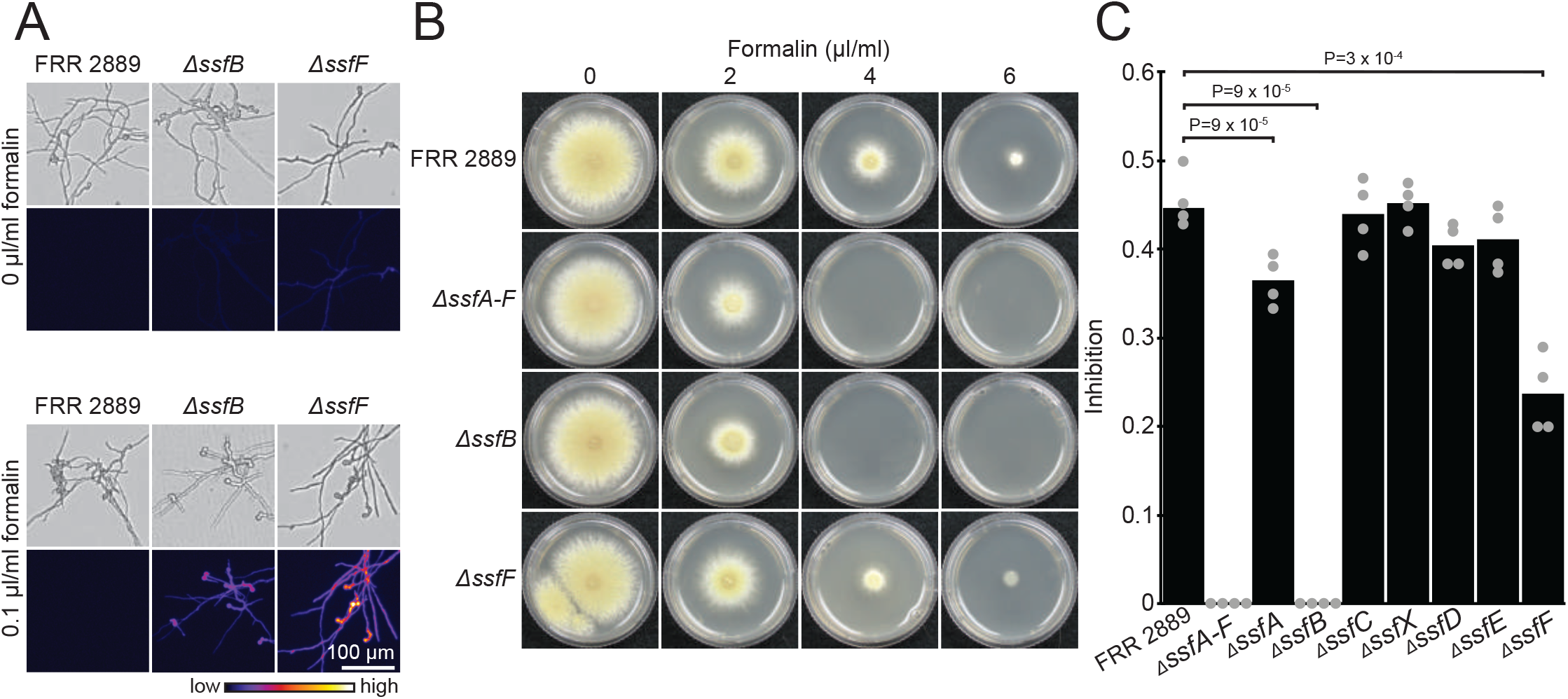
The *ssfA-F* subregion confers resistance to formaldehyde. **A)** GFP replacement strains indicated that some *ssf* genes, including *ssfB* and *ssfF* were strongly induced in response to formaldehyde (0.1 µl/ml formalin), a complete gene set is shown in Figure S6. **B&C**) Deletion of the entire *ssf* cluster, *ΔssfA-F*, resulted in decreased formaldehyde resistance, comparable to deletion of a single gene *ssfB*. The *ΔssfF* mutant showed a milder phenotype characterized by decreased growth rate at 4 µl/ml formalin. Plates were cultured at 30°C for 3 days

We then proceeded to examine the formaldehyde tolerance of a strain in which the entire *ssfA-F* region was replaced by GFP under the promoter of the *ssfA* gene (Δ*Χρ*+ Δ*ssfA-F*) (Figure 3B and C). The Δ*Χρ*+ Δ*ssfA-F* strain showed the expected increased sensitivity to formaldehyde with a lack of growth at 4 µl/ml formalin (Figure 3B and C). Consistent with the segregation analysis, tolerance in this strain remained higher than the *Χρ*-wildtype strains with the ability to grow at 2 µl/ml formalin being retained in the mutant. This is congruent with the hypothesis that a second unlinked locus in FRR 2889 and CBS 101075 contributes to increased formaldehyde tolerance in these two *Xρ+* wildtype strains. To confirm the presence of at least one additional formaldehyde resistance locus in the genome we crossed this *Χρ*+ Δ*ssfA-F* mutant to the *Χρ*-wild type CBS 144490. Among 19 progeny, 14 were able to grow on 2 µl/ml formalin and none at 4 µl/ml formalin. Resistance to 2 µl/ml formalin did not co-segregate with the presence of *Χρ*+, which was determined by hygromycin resistance that was introduced at the same time as GFP in order to select transformants (Figure S7). We did not identify the genetic bases for the difference in background resistance between *Paecilomyces* strains.

Having shown that the *ssfA-F* subregion was responsible for formaldehyde tolerance, we next investigated the individual contributions of each *ssf* gene (Figure 3B and C, S8). Examination of the formaldehyde resistance of the single gene GFP replacement mutants revealed greatly reduced growth in the *ssfB* mutant with a complete absence of growth at 4 µl/ml formalin. This suggested that *ssfB* is a major contributor to the increased formaldehyde resistance associated with the *ssfA-F* subregion, despite this gene having no clear role in formaldehyde tolerance based on characterised protein homologs. The biological role of *ssfB*, a disulphide oxidoreductase, is unclear; however one group of disulphide reductases, namely glutathione reductases, reduce glutathione disulphide (GSSG) to reduced glutathione (GSH) which maintain the oxidative balance of the cell. The reaction of glutathione and formaldehyde has been shown to alter the GSH:GGSG ratio and trigger redox stress in human cells and nematodes [58, 59]. As such a role for *ssfB* in maintaining reduced glutathione or redox balance more generally is possible. The identification of this enzyme is a promising lead for further biochemical investigation. *ssfF* had a more subtle effect on formaldehyde tolerance as assessed by radial growth rate at 4 µl/ml formalin. SsfF shows similarity to the formaldehyde-activating enzyme (Gfa) from the bacteria *Paracoccus* which is believed to increase the rate of reaction between formaldehyde and glutathione to form *S*-hydroxymethylglutathione [38]; however a more recent study suggests that Gfa does not directly catalyse this reaction [60]. This study is the first time that a phenotype has been linked to this enzyme in any organism. Complementation of the *ssfB* and *ssfF* mutants with wildtype copies of the genes restored formaldehyde resistance to near-wildtype levels (Figure S9). Deletion of the remaining *ssf* genes did not have a major impact of formaldehyde resistance under the conditions tested (Figure 3C).

### *ssfD* and *ssfF* are functionally redundant with their paralogs in the host genome

In addition to the *ssf* genes within *Χρ*, *Paecilomyces* genomes contain additional genes with putative roles in formaldehyde resistance which are distributed across presumably non-mobile regions of the genome. For example, the *P. variotii* genome has homologs of *ssfA*, *ssfC*, *ssfD*, *ssfE* and *ssfF*, but not *ssfB* and *ssfX* (Figure 2A and B). These “host paralogs” appear to be vertically inherited, as their phylogenetic relationships within the *Paecilomyces* genus match established species relationships (Figure 2A). The host paralogs form a distinct monophyletic group, to the exclusion of the clade of *Starship-*related *ssf* homologs. We hypothesized that some of the *ssf* genes might be partially or fully redundant with their corresponding host paralogs. Although the single gene KO that produced the strongest phenotype was *ssfB*, this gene lacks a clear paralog in *P. variotii*. Therefore, we chose to test this hypothesis by deleting the paralogs of *ssfD* and *ssfF* which we named *psfD* and *psfF* respectively (paralog of *ssf*). We chose *ssfD* because this gene has a putative role in the conversion of *S*-formylglutathione to formate + glutathione (Figure 1B). We anticipated that an *ssfD/psfD* double mutant would be highly impaired in formaldehyde resistance. Single deletions of *ssfD* or *psfD* had minimal impact on formaldehyde resistance. However an *ΔssfB/psfD* double mutant had increased sensitivity to formaldehyde with minimal growth even on 0.5 µl/ml formalin (Figure 4C & E). This suggests these two paralogs are functionally redundant and do not contribute additively to formaldehyde resistance under test conditions.

**Figure 4:**
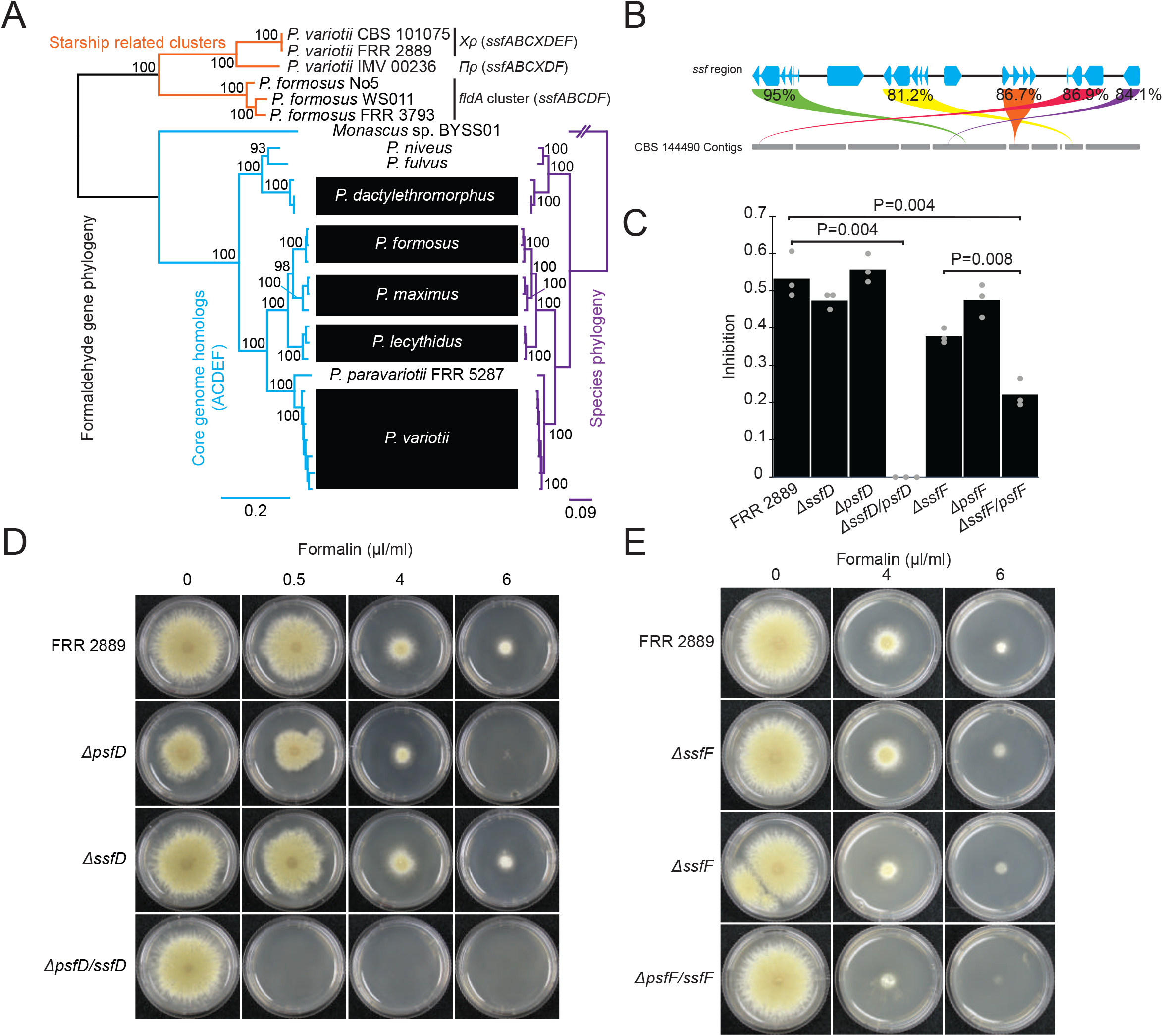
Some *ssf* genes are at least partially functionally redundant with the corresponding host homologs. **A)** Phylogeny of formaldehyde resistance genes (concatenated gene regions) in *Paecilomyces* strains compared to species relationships. Tree was generated in IQ-TREE with branch supports calculated from 10000 ultrafast bootstraps [61, 62]. The species phylogeny is taken from Urquhart et al. [45]. **B)** Similarity (BLOSUM62 [63]) between the Ssf proteins and their closest paralog in the *Χρ*-strain CBS 144490. **C&D)** Growth rate of *ΔssfD*, *ΔpsfD* and the *ΔssfD/psfD* double mutant on formaldehyde-containing media. **D&E)** Growth rate of the *ΔssfF*, *ΔpsfF* and the *ΔssfF/psfF* double mutant on formaldehyde containing media. Plates were cultured at 30°C for 3 days.

On the other hand, we chose gene *ssfF* based on its biological novelty as it has only been examined in prokaryotes and has never been experimentally linked to formaldehyde metabolism in eukaryotes. An *ΔssfF/psfF* double mutant showed decreased growth relative to wildtype on 4 µl/mL formalin plates while the single gene knockouts showed either no (*psfF*) or modest (*ssfF*) reduction in growth rate (Figure 4C & F). These results again suggest functional redundancy between host and *Starship* paralogs under the examined experimental conditions. The less dramatic effect of *ssfF/psfF* deletion compared to *ssfD*/*psfD* deletion is consistent with the fact that the reaction of formaldehyde and glutathione can be a spontaneous process which does not depend upon enzymatic activity [38].

### The *ssf* regions carried as *Starship* cargo illustrate birth-and-death processes with evidence of inter-*Starship* gene exchange and eventual incorporation of cargo into the host genome

To better understand the evolutionary history of the *ssf* cluster, we searched 2,899 publicly available fungal genomes sampled from across the fungal tree of life (Table S1), and generated individual phylogenies of *ssfB, ssfD, ssfX* and *ssfF*. All of the gene trees exhibit topologies that are incongruent with species relationships (Figure 5A; Figure S10-13; Supporting Data). We used these data to search for genomic neighborhoods containing clusters of *ssf* genes defined by having hits to at least 3 different genes within 25,000 bp distance of each other. There are remarkably few instances that fit these criteria across the fungal kingdom, consistent with these genes generally being unclustered (Table S2). We used the evolutionary history of *ssfB* as a proxy to understand the history of the *ssf* cluster and its associated phenotypes, as it makes the greatest contribution to formaldehyde resistance and is consistently present across all *Starship*-associated *ssf* clusters. All *ssfB* sequences form a supported monophyletic clade with the *Τυ* sequences sister to the other sequences from the other three *Starships*, consistent with the lower pairwise similarity of its *ssf* cluster to the others (Figure 1A). *Mι* and *Πρ* sequences form a single clade in agreement with a very recent exchange of the *ssf* cluster between them. Intriguingly, *Χρ* sequences are part of a clade with clusters that do not directly appear to be *Starship* associated. However, some species, such as *A. nomiae* and *A. bombycis*, have what appear to be degraded captain sequences nearby, indicating that they likely represent derelict *Starships*. Furthermore, a fragmented cluster from *A. flavus* that is sister to *A. nomius* shows extended homology with the *A. nomius* cluster, indicating that this region is likely also *Starship* derived (Figure 5A). Thus, we can infer that all *ssf* clusters have a single origin that coincides with their association to *Starships*, but that some clusters and cluster fragments have become integrated to the host background.

**Figure 5:**
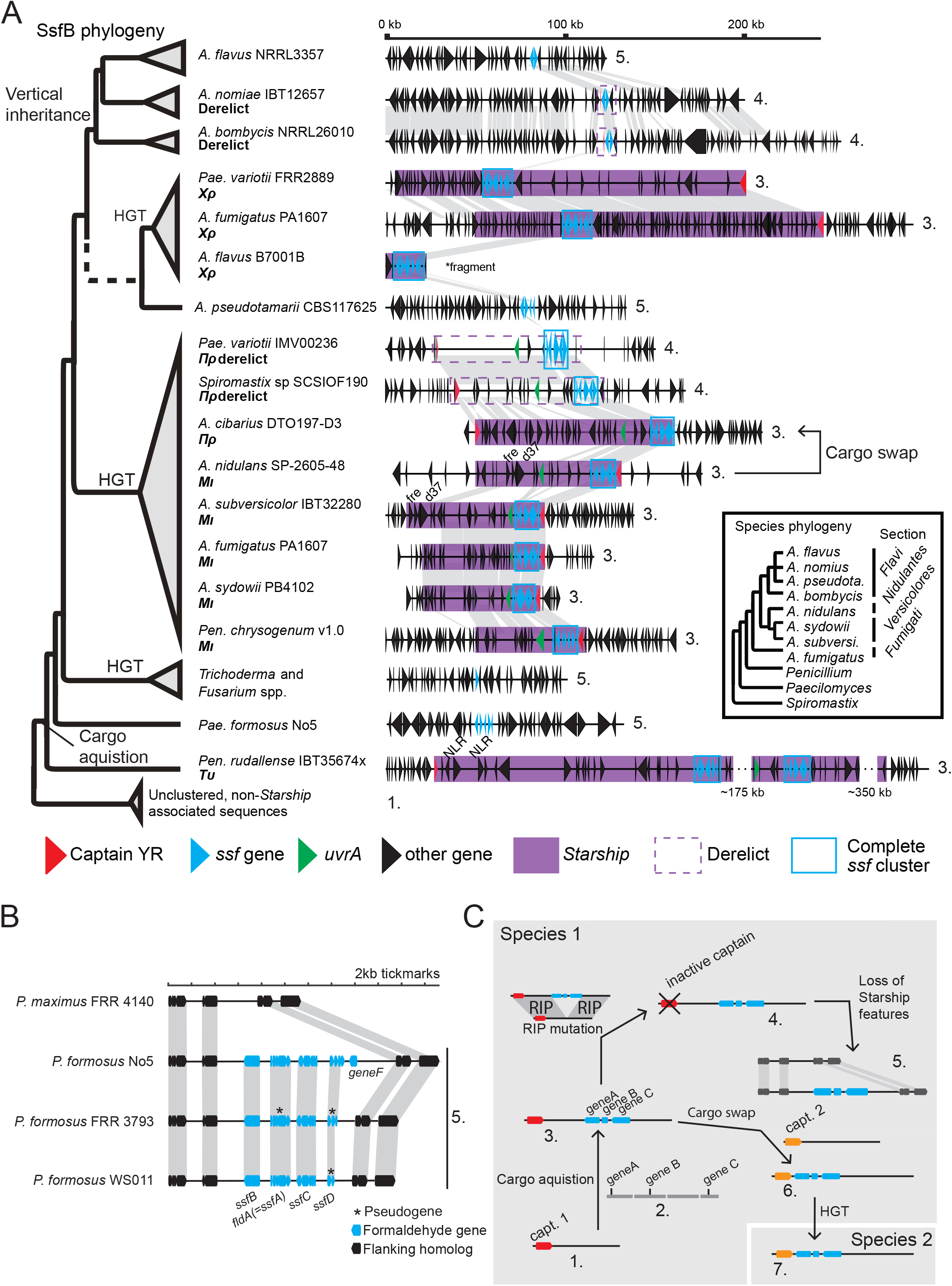
*ssf* clusters have moved both between *Starships* and into normal genomic regions. **A)** An SsfB protein phylogeny with corresponding *ssf*-cluster containing regions illustrated. The dashed line indicates a poorly supported branch (0.1% SH-aLRT and 57% UFboot support). “Cargo acquisition” refers to the original acquisition of the *ssf* cluster genes by a *Starship*.”Vertical inheritance” refers to a loss of *Starship* features followed by vertical inheritance of the ssf cluster. Numbers on the right hand side correspond to stages in the *Starship* lifecycle shown in part C. **B)** An *ssf* cluster in *P. formosus* which is homologous to the ssf cluster but is not associated with *Starship* features appears to represent an insertion into the genome which is degraded to varying extents in three different *P. formosus* strains. “5.” refers to step 5 in part C which is the step in the *Starship* lifecycle which we believe this cluster represents. **C)** Diagram illustrating the proposed “lifecycle” of a *Starship*. 1. An ancestral *Starship* not containing the cluster. 2. Unlinked genes within the genome. 3. A *Starship* in which the genes have become linked and and are now present as a cluster. 4. A derelict *Starship* which has been mutated and is now immobilized. RIP mutation refers to repeat induced point mutation. 5. Genes originally carried into the genome by a *Starship* now appear as an indel relative to a sister species. 6. A cargo swapping between *Starships* has moved the gene cluster into a different *Starship*. 7. Acquisition of the gene cluster provides an adaptive advantage to the strain carrying the *Starship* which promotes horizontal transfer into a different fungal species. An HGT event of the cluster-carrying *Starship* (3) could also occur.

The *ssf* clusters in *Mι* and *Πρ* are very closely related and highly similar (>98% nucleotide identity for each *ssf* gene), despite the extremely distant relationships between their respective captains (∼16% amino acid identity, Figure 1). Furthermore, these two *Starships* share a significant amount of common gene content beyond the formaldehyde resistance genes, including the *uvrA* gene [29] (Figure 5A), we previously observed that the *uvrA* gene is a putative DNA repair protein horizontally transferred into fungi from bacteria which is restricted to a small number of fungal species [29]. The striking similarity of cargo between two *Starships* from entirely different families strongly suggests that an inter-*Starship* recombination event has facilitated the de novo acquisition of the *ssf* cluster by one of these elements. Inferring the direction of HGT events is challenging because the phylogenies of captain and cargo genes often do not match the species tree at multiple internal nodes. The extended captain YR phylogeny presented in Figure 1C, which incorporates captains from *Starships* that are closely related to those examined here [33], suggests that both *Starships* originated in *Aspergillus*, as they are most closely related to other *Aspergillus*-associated *Starships* and were subsequently transferred to *Paecilomyces, Penicillium*, or *Spiromastigoides*.

We searched for additional evidence of HGT directionality by searching for cases where a smaller, species-specific TE may have jumped into the *Starship* prior to its transfer. In both *Spiromastigoides* and *Paecilomyces*, *Πρ* has a nested LINE element, but at unique insertion points. This element occurs only a single time in each genome, but is multicopy within the species *A. sydowii*, suggesting that it has been associated with *A. sydowii* for a longer period of time. Furthermore, *Spiromastigoides* has a second element belonging to the Prometheus-family, that shows high similarity to *Starship Πρ*, but lacks the *ssf* cluster. This *Starship* is nearly identical to one from *A. sydowii* CBS 593.65, and again possesses a nested copy of the same LINE element. Together, this evidence strongly supports an origin of *Πρ* in *A. sydowii* or a close relative with subsequent transfers to *Spiromastigoides* and *Paecilomyces*. As *A. sydowii* also possesses an *ssf*-carrying haplotype of *Mι*, this implies that *Πρ* acquired the cluster from *Mι*. Additionally, the non-*Starship* associated *ssf* cluster fragment in *A. nomius* and *A. bombycis* has some nucleotide similarity to the start of *Mι*, and so it may also be the case that *Mι* was the original donor of the *ssf* cluster to *Χρ* as well. As all *Starships* show evidence of HGT coincident with acquiring the *ssf* cluster, we infer that recurrent exchange of adaptive cargo genes between *Starships*, promotes subsequent transfer to other lineages. The divergence of *Τυ* relative to the other *ssf* clusters indicates that this process of repeated HGT has been occurring for extended amounts of evolutionary time, significantly impacting the evolution of these fungi.

Among the *ssf* clusters identified (Figure 5A) there is a spectrum of elements representing the stages in the “life cycle” of a *Starship* (Figure 5D). Most straightforwardly, there are some ssf clusters nested within intact *Starships* such as the *Χρ* elements in *Paecilomyces variotii* and *Aspergillus fumigatus*. In other cases, the ssf clusters are found within *Starships* that are clearly mutated and no longer active. The *Spiromastix Πρ* in *P. variotii* IMV00236 and *Spiromastix sp*. SCSIOF190 are heavily affected by repeat induced point mutation (RIP). We previously showed experimentally that if the starship *Hφ* is duplicated within the genome it can be heavily mutated via RIP in *Paecilomyces variotii* [29]. Interestingly, the RIP mutation was not present within the *ssf* region suggesting that the RIP occurred as a result of the presence of a second *Πρ* within the genome prior to the sexual cycle (Figure S14). Similarly, the *A. nomius* and *A. bombycis ssf* clusters are associated with a pseudogenised captain YR gene. Some other clusters show no *Starship* features, yet are sister to *Starship*-associated sequences. We believe that such clusters, such as those in *A. flavus*, *A. pseudotamarii*, *Trichoderma*, *Fusarium* and *Paecilomyces formosus*, represent former *Starship* cargo that has been integrated into the host genome. For example, we examined the cluster in *P. formosus*, which has no evidence of associated *Starship* sequence and is divergent from all other clusters. It is within this region that the *ssfA* homolog *fldA,* which was previously purified from *P. formosus* number 5, is found [39]. The genomic region is an insertion relative to the sister species *P. maximus*. In each of three available *P. formosus* genomes the region shows different degrees of truncation and pseudogenization suggesting that the region is actively being degraded. Of note, *ssfB* appears to be universally maintained, while other genes are pseudogenized or deleted (Figure 5B). Given the observation that these genes have redundant functions with homologs in the genome of *P. variotii*, it suggests that selection is primarily maintaining genes which uniquely contribute to formaldehyde resistance, tailoring the cluster to a specific fungal genomic landscape. Congruently, we see that the clade containing *A. flavus* sequences that are sister to *A. nomius* has only maintained *ssfA* and *ssfB*. Another piece of evidence that the *ssf* cluster in *P. formosus* is *Starship* derived, is that genes *uvrA* and *ssfX* are present in the genome, though located elsewhere. The presence of *uvrA* is particularly compelling as this gene appears to have originally been acquired through an HGT event from bacteria, and is only present in fungi that possess the *ssf* gene cluster (being found in the Starships *Πρ*, *Mι* and *Τυ*) with the exception of *Phaeoacremonium* (Figure S15). The fact that horizontally and vertically inherited *Starship* cargo may eventually incorporate into the host genome is likely to lead to an underestimation of the importance of *Starships* in fungal evolution as such regions are difficult to recognise. Crucially, we have no evidence that the *ssf* cluster must be uniquely associated with *Starships* or somehow increase the rate of HGT events. Ergo, the dynamics uncovered herein should be applicable to any fungal phenotype that can be controlled by one or a few genes. Together with the vast array of functionality encoded by *Starship* cargo this implies that these processes are paramount to the evolution of fungi and their rapid adaptation to novel niches.

## Discussion

### The *ssf* cluster supports the hypothesis that *Starship* HGT and cargo swapping represent a powerful force in fungal adaptation and elevates the role of HGT in eukaryotes

We now have a number of examples of HGT events from a wide diversity of eukaryotes including fungi [14–17, 20, 22, 23, 30, 64, 65]. However, much about the role of HGT in eukaryotic biology remains unclear. On one hand, HGT among eukaryotes might be restricted to isolated events, which while important in specific cases, are not major drivers of evolutionary change. The alternative possibility is that there are active mechanisms of HGT in eukaryotes, similar to those in prokaryotes, and that these processes mobilize genetic information between species at relatively high rates, repeatedly driving adaptation. Examining the *ssf* cluster demonstrates that *Starships* repeatedly facilitate the evolution of an adaptive trait through inter-element cargo swapping and HGT, supporting a model in which HGT plays a more dominant role in fungal evolution. We have found the cluster to be present in numerous fungal species across different genera as a result of HGT (Figure 1). Moreover, we have found that at least four phylogenetically distant *Starships* have acquired and then mobilized the *ssf* cluster between fungal species following the swapping of cargo content between different *Starships*. There is no precedent among eukaryotes for a genetic region so repeatedly associated with horizontal gene transfer with at least nine HGT events inferred. Given the relatively few publicly available genomes compared to extant fungal diversity, this figure likely massively understates the true number of HGTs of the *ssf* cluster driven by *Starships*.

### Other gene regions appear to follow the same evolutionary pattern as *ssf*

While the *ssf* gene cluster provides the most illustrative case study, two other examples of published *Starships* can now be interpreted to fit a similar evolutionary pattern [30, 31]. The first is microbial host range driven by the *Starship* Horizon in pathogens of wheat [31]. The *ToxA* gene encodes a proteinaceous host-specific toxin (TOXA), which enables pathogenicity on wheat cultivars with a particular susceptibility gene *Tsn1* [66]. Since the mid 2000s it has been clear that multiple wheat pathogens have utilised the common strategy of using the TOXA protein to infect susceptible wheat. These include *Pyrenophora tritici*-*repentis* [67, 68], *Parastagonospora nodorum* [19] and *Bipolaris sorokiniana* [69]. The acquisition of *ToxA* by *P. tritici-repentis* likely occurred around 1940 and resulted in the emergence of this species as a new agent of disease in wheat [19]. McDonald *et al*. showed that in each of these species *ToxA* was located within the 14 kb transposon ToxhAT, which in each species was internally embedded within a larger insertion relative to strains lacking *ToxA*. Due to the high level of sequence similarity within ToxhAT, HGT between these pathogens appeared to be the only reasonable hypothesis to explain its presence in multiple species [19]. It was initially proposed that ToxhAT was responsible for mediating the HGT event [31, 70]. However, it has been recently discovered that the insertion within which ToxhAT is found is in fact a *Starship* named *Horizon* in both *Pyrenophora tritici*-*repentis* and *Parastagonospora nodorum*; and in an unrelated unpublished *Starship* named *Sanctuary* in *B. sorokiniana*, raising the alternative hypothesis of *Starship* mediated HGT [31, 71]. In light of the evolutionary dynamics and cargo swapping of *ssf*, which in the case of *ToxA* may be aided by the mobility of ToxhAT, we strongly favor the *Starship*-mediated hypothesis.

The second example of *Starship* driven adaptation is the evolution of metal resistance in Eurotiales species through acquisition of Hephaestus-family *Starships*. Numerous Hephaestus-family *Starships* have independently acquired genes that encode resistance to various metals based on bioinformatic analysis of gene content. However, the best studied Hephaestus-family *Starship*, the type element *Hφ,* found in *P. variotii,* confers resistance to at least five different metal ions zinc, copper, arsenic, cadmium and lead mediated by four different genes with cadmium and lead resistance the result of a single transporter gene [30]. As with *Starships* containing the *ssf* gene cluster, we found evidence for interactions between *Starships*. Specifically, the genes required for arsenic resistance appear to have been introduced into *Hφ* as result of a nesting event [30]. Initially we identified a signal of HGT of this element between *P. variotii* and *Penicillium fuscoglucum* [30]. Since then we have observed additional HGT events between *P. variotii* and the *P. lecythidis* strain MCCF 102. This was a particularly striking example as the two copies of *Hφ* differ by a single nucleotide across 86 kb[29]. The sister species of *P. variotii*, *P. paravariotii* (only recently delimited from *P. variotii*) also contains apparent HGT mediated acquisition of the element [45]. Finally, examination of the recently released *Penicillium chermesium* IBT 19713 genome reveals yet another HGT of this element (Figure S4) [54]. We thus have evidence for at least four HGT events of this *Starship* just in publicly available genomes.

In addition to these two examples which are explicitly described as *Starships*, there are already a number of HGT events described in the literature which are likely to have been mediated by *Starships*. For example, we have noted that several of the HGT regions reported among cheese making fungi are *Starships* [28, 29]. Similarly, a number of horizontally transferred regions in *Penicillium* fungi used to make salami were reported to contain the Captain YR enzymes with the DUF3435 domain which suggests that these regions of HGT are also *Starships* [72]. The SORBUS gene cluster of *P. roqueforti* shows presence/absence polymorphism among strains and increases resistance to sorbic acid. This region is suspected to have been introduced into the *P. roqueforti* genome via HGT and contains a DUF3723 which is a characteristic feature of *Starships* [53]. As such, the supposition that the *Starhsips* operate as active mechanisms of HGT to mediate recurrent, rapid adaptation beyond just formaldehyde resistance has tangible avenues for future investigation.

#### The selective advantage conferred by the cargo increases the probability of a *Starship* persisting following HGT

It is noteworthy that in the three examples of *Starship*-mediated HGT that have been best studied, namely *ToxA*, metal resistance, and now formaldehyde resistance, in each case HGT has occurred multiple times. The prevalence of HGT is particularly interesting given that the majority of *Starship* evolution appears to follow a vertical pattern of inheritance [28]. We hypothesize that *Starships* which gain host-beneficial cargo are primed for rapid dissemination through horizontal transfer events because these elements provide a competitive advantage to the recipient genome and it is thus more likely that this recipient genome will pass its genes to the next generation into the wider gene pool. In the absence of selective pressure, horizontally transferred elements are expected to be lost via genetic drift in most cases just as with any other neutral or nearly neutral mutation. A feature shared by both the *ssf* gene cluster and toxA is that they are found within multiple distantly related *Starships*, suggesting that the reshuffling and swapping of adaptive cargo is common. Inter-mobile element gene shuffling is frequently observed in bacterial plasmids and integrative and conjugative elements and is a critical process generating not only new combinations of functional modules but in promoting the maintenance and horizontal spread of ecologically and economically important phenotypes like antibiotic resistance and nitrogen fixation [73–75]. It remains to be seen how often *Starships* exchange cargo compared with their bacterial analogs and what environments or genomic contexts might promote reshuffling.

### Formaldehyde is a ubiquitous organic compound found in a wide range of environments making it difficult to understand in which environmental contexts the *ssf* cluster confers its adaptive advantage

The high level of identity between the *Starship* regions as shown in Figure 1 beyond indicating the occurrence of HGT, implies that these events must have occurred in the recent past. It is thus interesting to consider the possible environmental factors driving the proliferation of this cluster. Formaldehyde is a ubiquitous organic compound and an important metabolite in biological systems (reviewed by [76]). Formaldehyde is found in high concentration in some fruit and vegetables which is one possible context in which saprotrophic members of the order Eurotiales might encounter formaldehyde [77]. For example, it is produced by the activity of alcohol dehydrogenases on methanol [78]. Formaldehyde is also an important anthropogenic pollutant, particularly of indoor air [79, 80]. Formaldehyde is toxic because it is highly reactive [81]. Living organisms therefore possess conserved mechanisms to limit this toxicity. In particular the glutathione-dependent formaldehyde detoxification pathway is utilized by a wide range of organisms including bacteria, fungi and other eukaryotes [34–37]. Several studies have isolated formaldehyde-resistant fungal strains from formaldehyde-contaminated environments [40, 82–85][86], suggesting that formaldehyde resistance might be rapidly evolving in response to environmental formaldehyde exposure. In a direct parallel to formaldehyde resistance mediated by *Χρ*, a conjugative plasmid has been demonstrated to transfer formaldehyde resistance between bacteria [87] suggesting that common selective pressures may act across both domains of life. However, given the wide range of sources of environmental formaldehyde we cannot know in which environmental context the *ssf* cluster confers its selective advantage. Indeed since formaldehyde is an intermediate in many biological pathways the advantage of the *ssf* cluster may even be conferred as a result of metabolizing endogenously produced formaldehyde, rather than detoxification of external sources of formaldehyde. Recently, the *Starship Swordfish* linked environmental data to *Starship* presence/absence across over 1000 *Zymoseptoria* strains isolated from across the world and found an association between the *Starship* and various climatic variables [32]. Future similar studies sequencing Eurotiales fungi directly isolated from different environmental contexts may be similarly illuminating in understanding the adaptive advantage of the *ssf* cluster in ecosystems.

### Conclusion - through mediating HGT *Starships* are a powerful force in fungal adaptation which should not be underestimated

Given that the acquisition of host beneficial gene cargo primes *Starships* for HGT events the *Starships* hold a so far unique position in eukaryotic evolutionary biology as agents of repeated adaptation via horizontal gene transfer. We show here that *Starship*-mediated HGT occurs repeatedly as opposed to in singular isolated events, indicating that HGT carried out by *Starships* is a powerful force in fungal adaptation the implications of which we are just beginning to recognise.

As shown via examination of the *ssf* cluster, the expected “endpoint” in the lifecycle of a *Starship* is that the cargo genes are integrated into the normal host genome, identifying such regions in the absence of *Starship* features is difficult. For example, T-toxin virulence genes of *Cochliobolus heterostrophus* are located in a genomic region consistent with a degraded *Starship* but lacking conclusive evidence [88]. Given the abundance of *Starships* in fungal genomes and their proclivity for horizontal transfer we suggest that in all cases of fungal to fungal HGT, *Starship*-mediated transmission represents a plausible explanation. Giant transposable elements are found in other eukaryotes for example as Teratorn elements are found in teleost fish and *Maverick* elements are found in a wide range of eukaryotes from insects to ciliates [89, 90]. Given that the HGT of a *Maverick* transposon has recently been reported in animals [91], we suggest that similar processes may be observed in many eukaryotic phyla. Given the diversity of threats and opportunities imparted by fungi and the potential for *Starships* to spread a wide range of gene cargos, they have potential implications on medicine, biosecurity and conservation which warrant urgent investigation.

## Methods

### Strains

Five wildtype strains of *Paecilomyces variotii* were examined i.e. CBS 144490, CBS 101075 (*Χρ*+), FRR 1658, FRR 2889 (*Χρ*+) and FRR 3823 for formaldehyde resistance all of which have available whole genome sequencing data [30, 92]. Genetic manipulations were conducted in strain FRR 2889 because this strain contained a full-length *Χρ* region. *Aspergillus fumigatus* strains selected from a recent pan genome study [56] including the only available *Χρ+* strain C-1-80s-1 and 12 additional strains (all *Χρ*-) as controls, namely C-1-76s-1, L-2-15-4, B-1-70s-1, E-1-66s-1, E-1-25-10, K-1-8L-3, C-1-66L-2, L-2-7-2, A-2-49L-2, L-2-11-2, B-1-68s-1, K-1-48s-1.

### *Paecilomyces* sexual crosses

*Paecilomyces* crossing was conducted as described previously [92, 93]. Strains of opposite mating type were plated on opposite sides of 90 mm petri dishes containing potato dextrose agar (PDA) and incubated in darkness at 30°C. Once ascospores were observed (up to 6 weeks) sexual spores were transferred to water and heat shocked at 80°C for 10 minutes to kill any contaminating conidia. The heat-treated spores were then plated onto CV8 agar and allowed to germinate for two days. The presence of *Χρ* within the progeny was assessed via PCR with duplex PCR using primer pairs AUB662+AUB663 which amplified 153 bp of the empty site of CBS 144490 and primer pair AUB651+AUB653 which amplify 204 bp within *Χρ*.

### Formaldehyde sensitivity assay

Formaldehyde sensitivity was assessed using 10% cleared V8 agar pH 6 for *Paecilomyces variotii* and using PDA for *Aspergillus fumigatus* supplemented with formaldehyde in the form of formalin (37% formaldehyde in H_2_O, containing 10-15% methanol as stabilizer). Strains were grown on the formaldehyde-containing media for 3 days at 30°C before radial growth was assessed.

### Cloning

PCRs were conducted using Q5 High-Fidelity 2× Master according to the manufacturer’s directions and the products purified using the QIAquick gel extraction kit. Specific details of the PCRs required for each construct are given in Table S3 Templates were either plasmid DNA or genomic DNA extracted via a CTAB-based protocol from lyophilised mycelium [94]

Fragments for knockout/GFP replacement constructs were cloned into EcoRI/HindIII double digested plasmid PLAUB36 which is the *Agrobacterium* binary vector pPZP-201BK modified to include the 2 micron origin and the *URA3* selectable marker for cloning in yeast [29]. The cloning step was conducted using homologous recombination in yeast by transforming the digested vector and purified PCR products into strain BY4742 using a standard LiAc/PEG approach [95]. Plasmids were rescued from the yeast and transferred to *E. coli* using the Zymoprep Yeast Plasmid Miniprep kit.

Complementation constructs were constructed by cloning the wildtype gene into the EcoRV site of plasmid pMAI2 [96] using the NEBuilder HiFi DNA Assembly Cloning Kit according to the manufacturer’s directions.

**Table S3.**
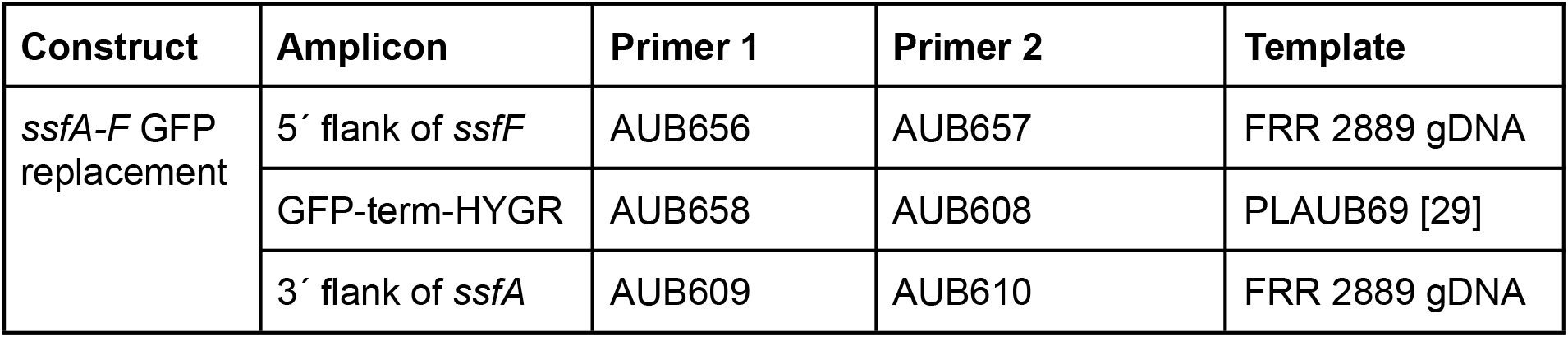
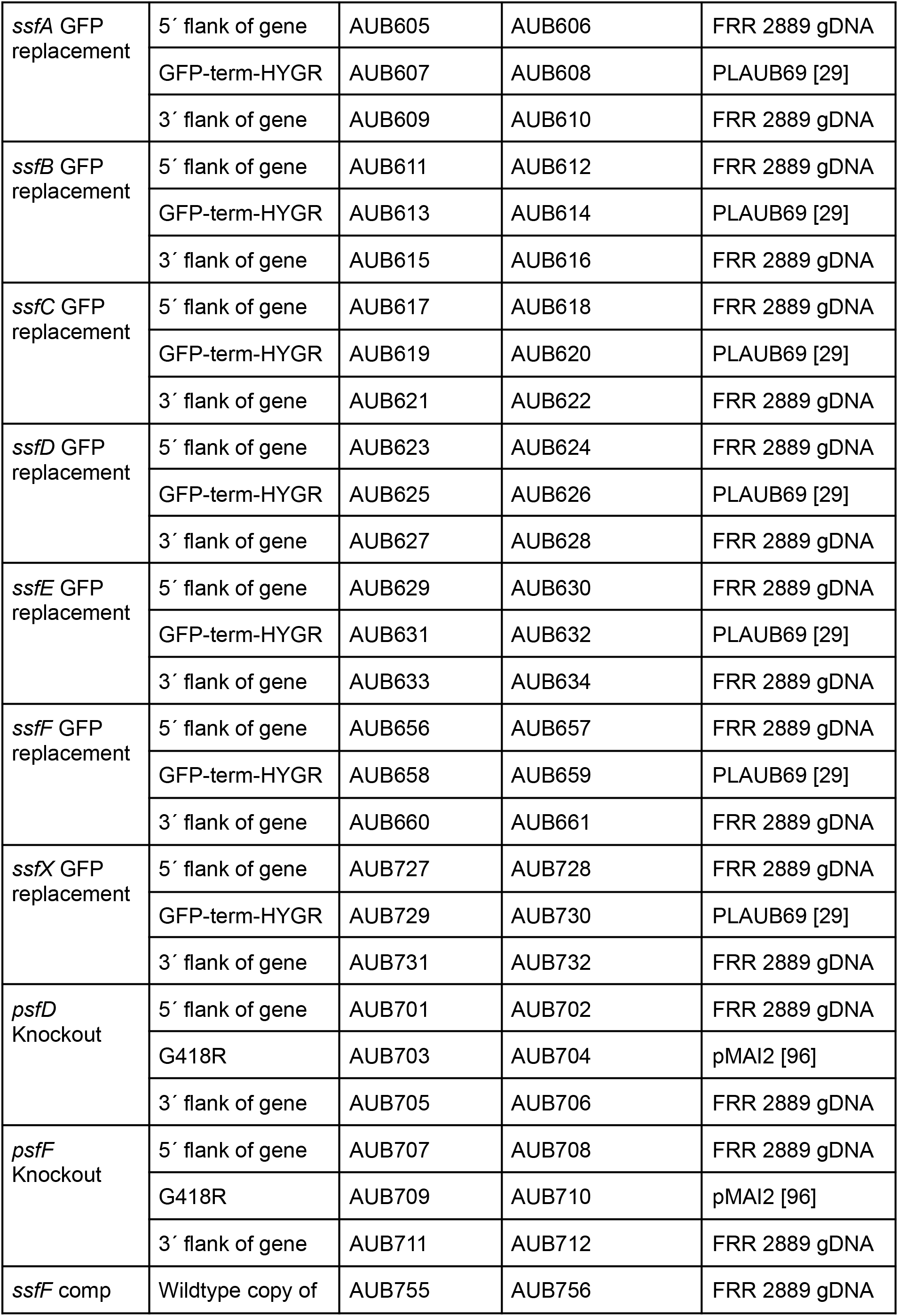
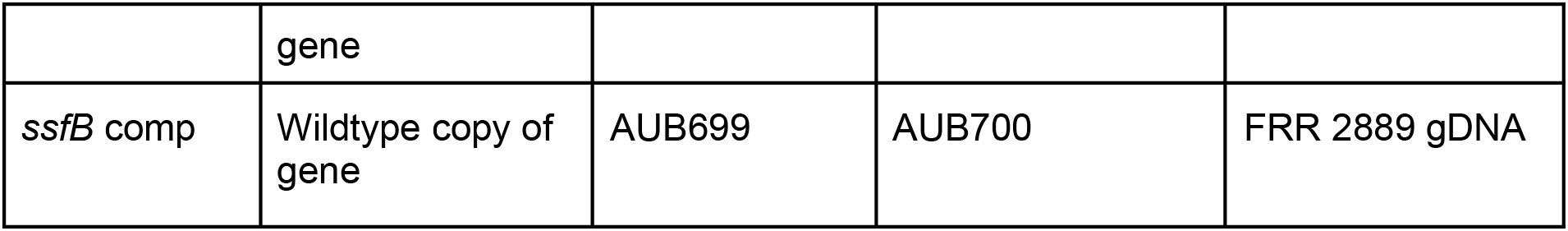

### Transformation

*Paecilomyces* was transformed as previously described [92]. Briefly, plasmid constructs were introduced into Agrobacterium strain EHA105 by electroporation. *Paecilomyces* was then co-cultured with the *Agrobacterium* on induction media containing acetosyringone before overlaying with selective media containing either hygromycin or G418 as appropriate for the construct. Transformant colonies were purified via single spore isolation before further analysis. In the case of gene knockouts, correct integration of the construct was assessed via PCR with the primers listed in Table S4. The “upstream” primer was paired with primer AUB696 in GFP, or AUB113 in the promoter driving the G418 resistance gene, to produce a product unique to a correctly integrated construct. Similarly, the “downstream” primer was paired with AUB114 in the terminator of both the hygromycin and G418 resistance genes to produce a product unique to a correctly integrated construct.

**Table S4.**
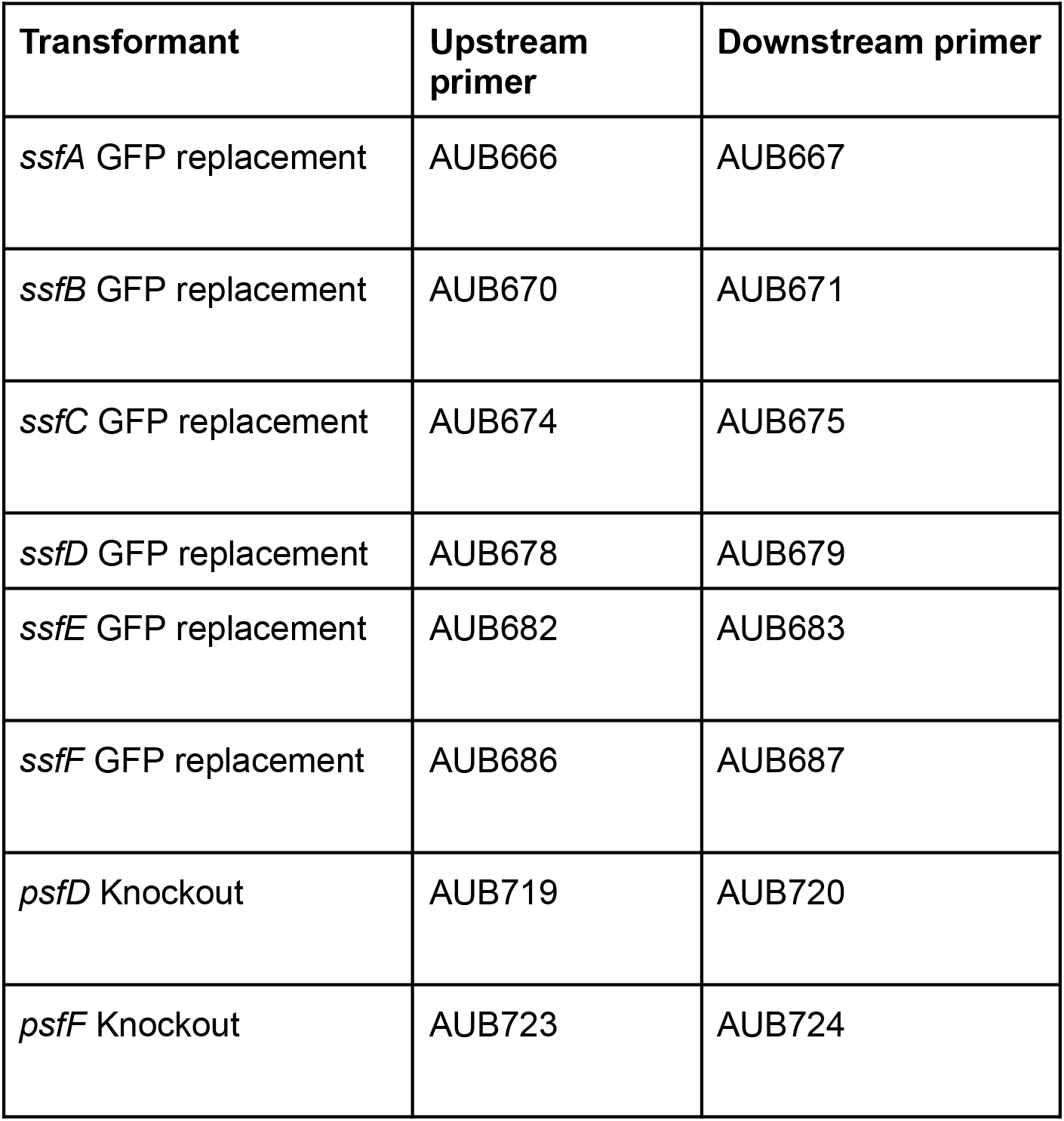
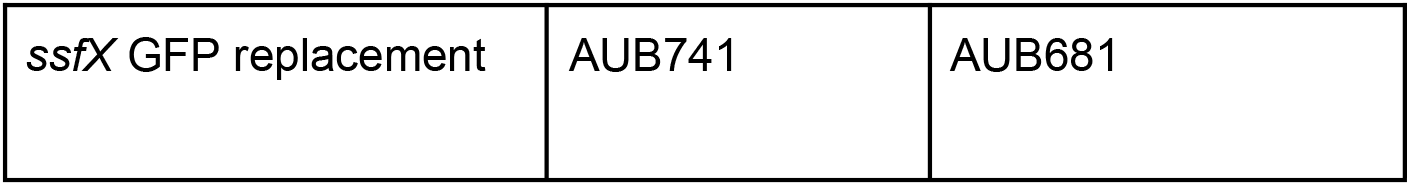

### GFP expression reporter

Spores were germinated overnight in PDB with or without 0.1 µl/mL formalin in still culture at 30 °C. Germlings were examined on microscope slides using a BioTek Cytation 1 imager. Settings were consistent across samples (4× objective GFP: LED intensity 5, integration time 350 ms, gain 0.6; Brightfield: LED intensity 5, integration time 38 ms, gain 0).

### Bioinformatic analysis

#### “Blast-all” inference of HGT

There are various methods by which HGT can be inferred. One approach is to use phylogenetic discordance i.e. to look for genes whose phylogenetic history differs significantly from expected species histories [98]. A weakness of phylogenies in the case of *Starships* is that these regions display presence/absence polymorphisms so we cannot place these genes into the context of a set of vertically inherited homologs. However, we do see that both captain YR and *ssf* cargo trees are highly discordant with expected species relationships. Here we have taken an alternative approach which we refer to as “BLAST-all”. In this method we take all the nucleotide gene sequences of one strain (predicted using Augustus implemented in Geneious Prime version 2023.0.4 with an *A. nidulans* gene model [97]) and use BLASTn to search these against the genomic scaffolds of another strain. We then take the top result for each gene and produce a scatter plot of identity vs alignment length. Genes which are more similar between two species will produce BLAST hits of greater length and higher identity. Genes that have recently transferred horizontally between species will be more similar than the rest of the genome, as they have diverged more recently compared with the rest of the genome. This phylogeny-agnostic approach of inferring HGT using sequence conservation in the context of whole genome comparisons is well suited for detecting large chunks of recently transferred DNA fragments, similar to the approach taken to infer HGT of *Maverick* transposons between nematode species [91]. An alternative explanation for the existence of conserved DNA between species is introgression, that is to say hybridisation between two species followed by repeated backcrossing until only the *Starship* remains. We do not think that this explanation is satisfying for at least two reasons. Firstly, as we described previously, the *Pegasus Starship* contains duplicated DNA and has horizontally transferred into *P. variotii* which is a species in which duplicated DNA is mutated during sexual crosses, thus the required backcrossing cannot have taken place. Secondly, while hybridisation events have been reported between different fungal species to our knowledge in all cases these occur between species within the same genus [99–101]. *Paecilomyces* and *Spiromastigoides* belong to separate orders. However, it remains possible that introgression plays a role in the transfer of the *ssf*-containing *Starships* between closely related species.

#### Sequence searches and phylogenetic tree building

We searched a database of 2,899 publicly available genomes for homologs to *ssf* cluster genes using BLASTp (e-value = 0.001) (Table S1). Due to the large amount of initial hits recovered for each gene, we iteratively built phylogenetic trees and selected clades of interest for further analysis based on their proximity to focal *ssf* genes from *Starship*-associated clusters. Each clade of interest was aligned using the E-INS-I method implemented in MAFFT version 7.490, which is recommended for sequences where several conserved motifs are embedded in long unalignable regions [102], and automatically trimmed using clipkit (--mode kpic-smart-gap) [103]. We built three maximum likelihood trees for each gene using IQ-TREE v2.0 with automated model selection, 1000 SH-aLRT tests and 1000 ultrafast rapid bootstraps (-m MFP -alrt 1000 -B 1000), and selected the one with the highest likelihood [61, 62].

A similar approach was used to build phylogenies specifically for *ssf* cluster genes found in *Paecilomyces.* Genes homologous to those found in the *ssf* cluster were identified via BLASTn searches against the same set of publicly available *Paecilomyces* genomes examined by Urquhart and Idnurm [45]. Each group of homologs was aligned using MAFFT version 7.490 [102], alignments were manually examined and trimmed to remove poorly aligned DNA. Alignments were then concatenated as follows. Those homologs clustered with other ssf genes were considered “core” homologs. The core homologs from each species were concatenated. The homologs found within a particular cluster were concatenated together. A tree was generated from the concatenated alignment using the default settings of IQ-TREE with branch supports calculated from 10000 ultrafast bootstraps [61, 62]. This phylogeny was compared to a species phylogeny taken directly from Urquhart and Idnurm [45].

#### Genomic neighborhood annotation

We searched for genomic neighborhoods containing clustered genes of interest using the complete set of BLASTp hits recovered for the *ssf* cluster genes from the 2,899 genome data as input to the command “starfish sketch” part of the starfish workflow [33]. Neighborhoods were defined as having hits to at least 3 different genes within 25,000 bp distance of each other. We extracted genes +/− 25,000 bp of these neighborhoods and included them as part of each neighborhood to better understand their genomic contexts (Table S2).

#### Determination of the presence of *Χρ* in *A. fumigatus* and *P. variotii* genomes

To confirm the presence of absence of *Χρ* in each strain Illumina sequencing reads were mapped to the FRR 2889 *Χρ* using inbuilt read mapper of Geneious Prime using the “low sensitivity/fastest” setting. Read depth across the *Χρ* region was assessed for presence/absence polymorphism.

#### Taxonomic determination of *P. rudallense* and *A. sydowi*

“*Pencillium sp.”* strain IBT 35674x [54] was identified as *A. rudallense* based on comparison of the *RPB2* sequence to those examined by Houbraken et al. [52]. Similarly, “*Aspergillus* sp.” PB4102 was taxonomically identified as *A. sydowii* based on a phylogenetic comparison of the *RPB2* sequence to those examined by Jurjevic et al [47]. In both cases the *RPB2* sequence of the strain of interest was extracted from the genome assembled using BLAST then aligned to the *RPB2* sequences of a set of taxonomically defined strains using MAFFT version 7.490 [102]. A tree was generated from the alignment using the default settings of IQ-TREE with branch supports calculated from 10000 ultrafast bootstraps [61, 62].

## Data availability

Scripts used in this study to generate figures and conduct comparative and statistical analyses, in addition to supporting data such as sequences, alignments and Newick tree files are publicly available from the following Figshare repository (DOI: 10.6084/m9.figshare.24552985).

