## Supplemental Figures and Tables S3 and S4 for "Gene acquisition by giant transposons primes eukaryotes for rapid evolution via horizontal gene transfer"

### Supplementary Figures

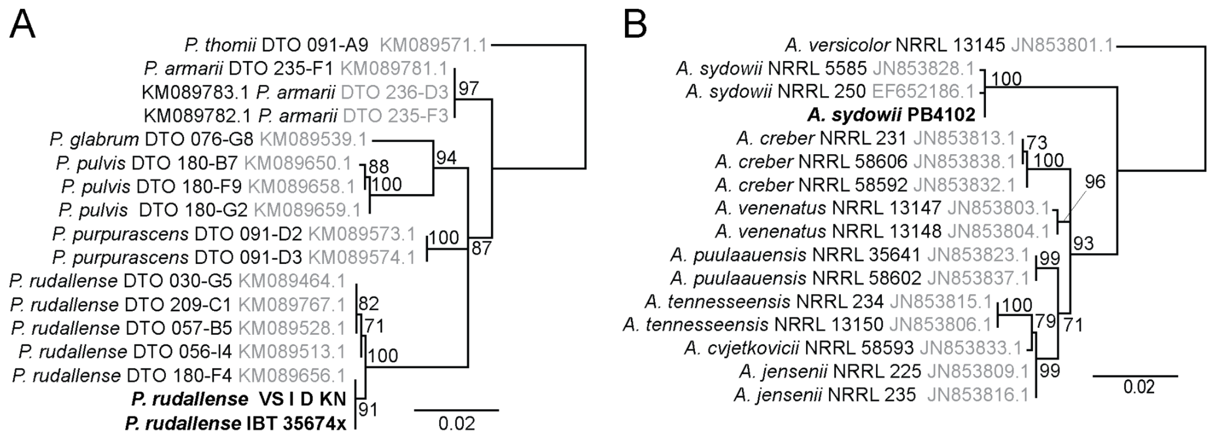


**Figure S1:** Taxonomic determination of *P. rudallense* and *A. sydowii* using the RPB2 nucleotide sequence. Trees generated in IQ-TREE. Ultrafast bootstraps from 1000 replicates indicated.

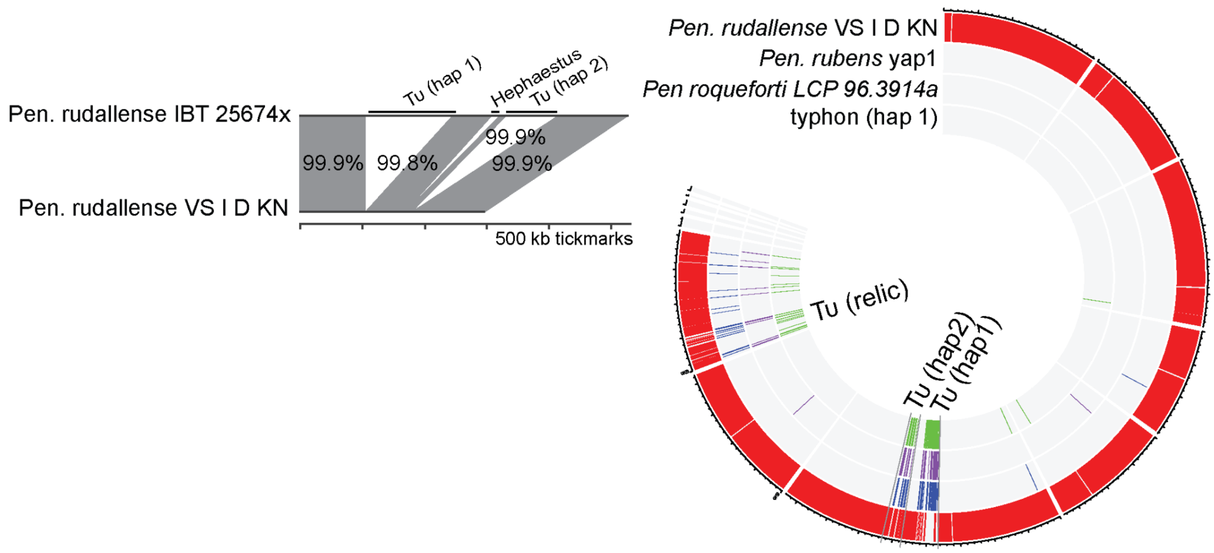


**Figure S2: A)** Alignment of the *P. rudallense* IBT 35674x contig containing *Tu* and a contig from *P. rudallense* VS I D KN containing the corresponding empty site. Three large insertions are present in *P. rudallense* IBT 35674x. Two of these are *Tu* Starships and one is an Hephaestus-family *Starship*. **B)** Distribution of *P. rudallense* IBT 35674x genes returning BLAST hits with length >100 bp and identity >97% from databases consisting of 3 genomes the conspecific strain VS I D KN. *P. rubens* YAP1, *P. roqueforti* strain LCP 96.3914a, or a database consisting of only the *Tu* haplotype 1 sequence.

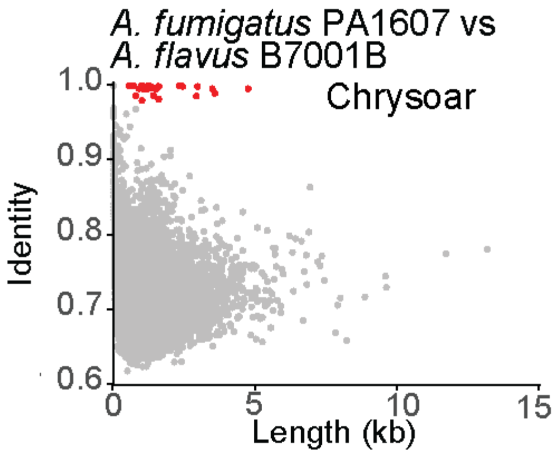


**Figure S3:** BLAST-all comparison of *A. fumigatus* PA1607 and *A. flavus* B7001B as per Figure 1D.

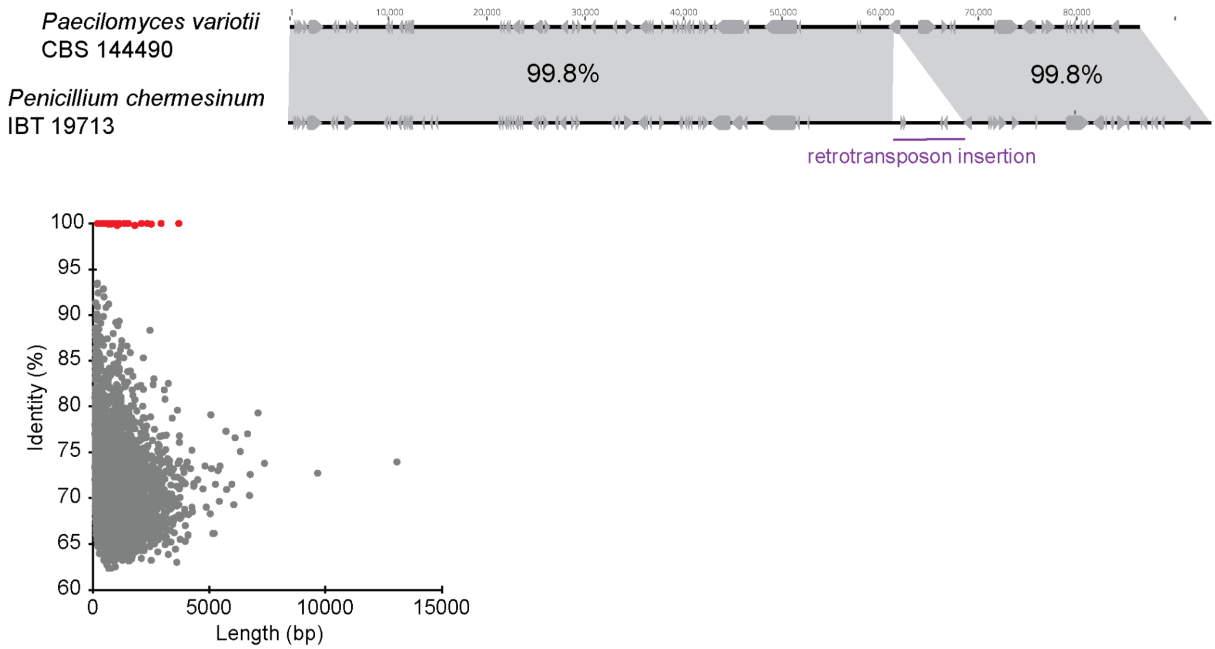


**Figure S4:** *Paecilomyces variotii* CBS 144490 and *Penicillium chermesinum* IBT 19713 contain near identical copies of the metal resistance *Hφ Starship* except for the insertion of a retrotransposon into *P. chermesinum.* The top panel shows the nucleotide alignment between the two copies and bottom panel the outcome of BLAST of all gene sequences, with the red dots being those found within *Hφ*.

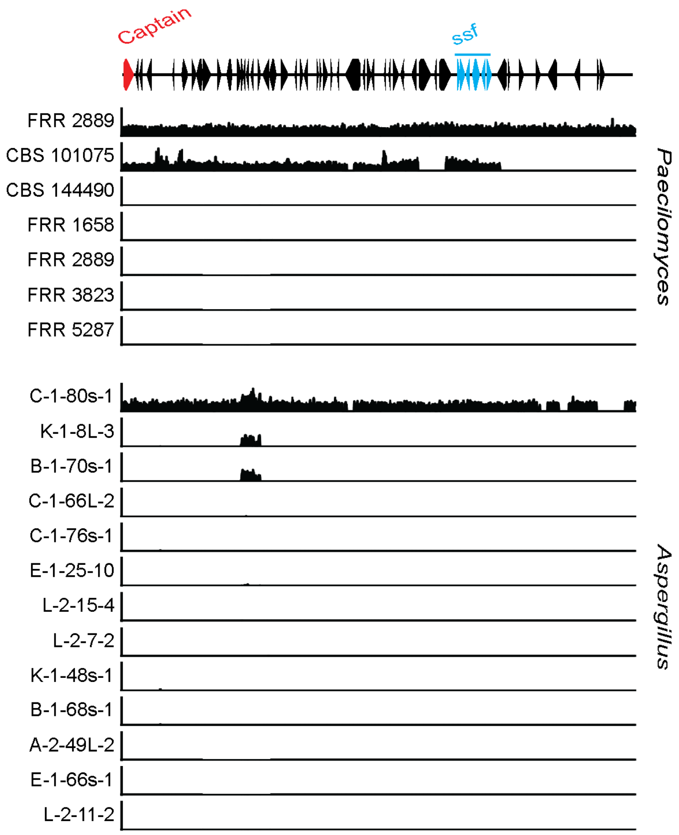


**Figure S5:** Illumina reads (5 million for each strain) mapped to the FRR 2889 *Xp* sequence. Y-axis represents read depth between 0 and 100× coverage.

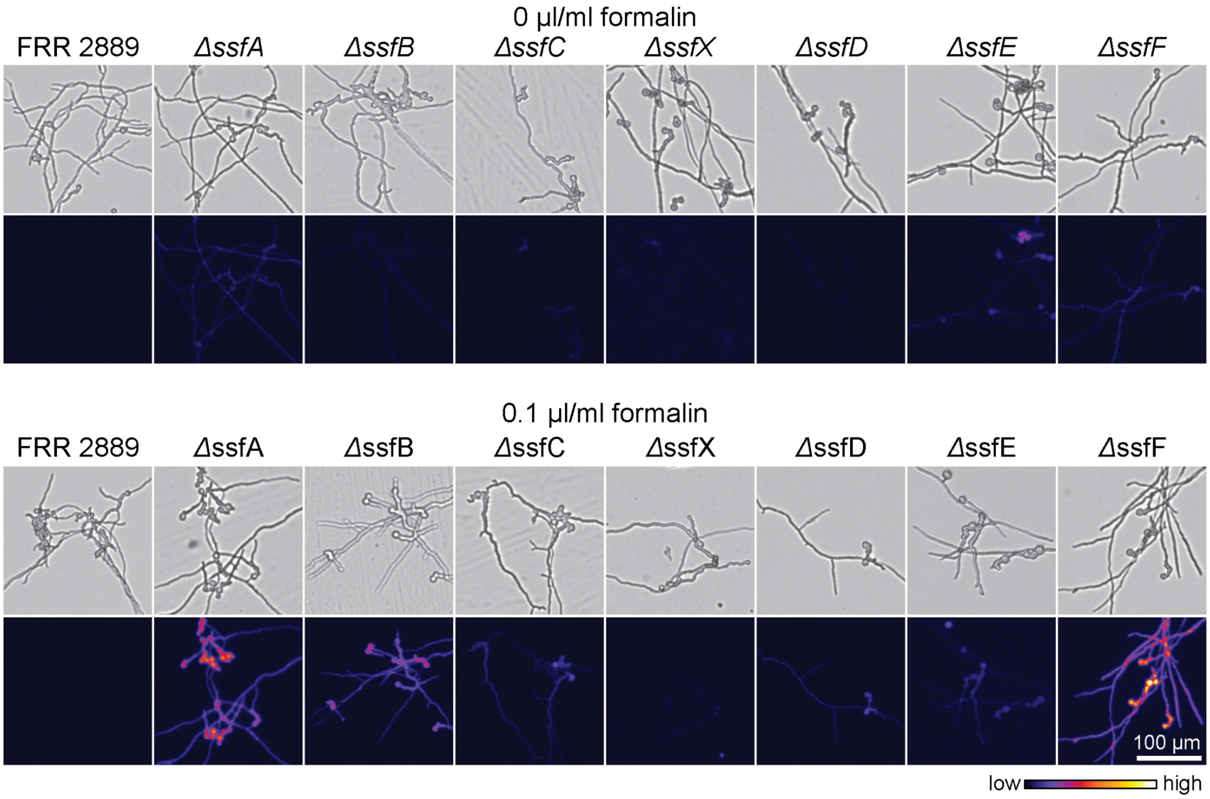


**Figure S6:** GFP fluorescence in wild type and GFP replacement mutants germinated overnight in potato dextrose broth with and without supplementation with 0.1 µl/ml formalin.

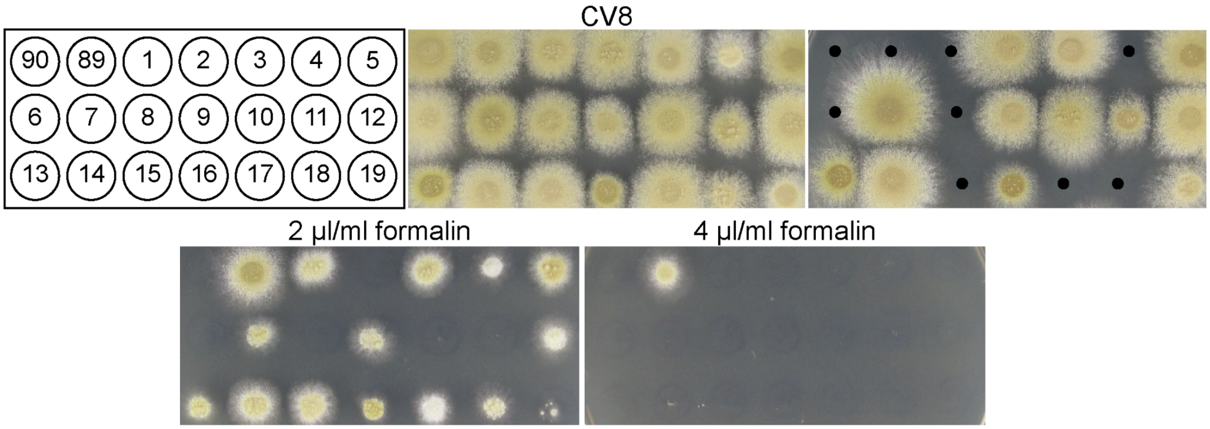


**Figure S7:** 19 progeny of a cross between CBS 144490 and FRR 2889 *ΔssfA-F* grown on cleared V8 agar (CV8), media supplemented with hygromycin (resistance is linked to *Xρ*), and media supplemented with 2 µl/ml or 4 µl/ml formalin. No progeny grow on 4 µl/ml formalin and on 2 µl/ml formalin resistance segregates independently to *Xρ*/hygromycin resistance. Black dots added to indicate lack of growth on hygromycin. The wildtype strains CBS 144490 ("90") and FRR 2889 ("89") are shown for comparison. Plates were incubated for 3 days at 30°C.

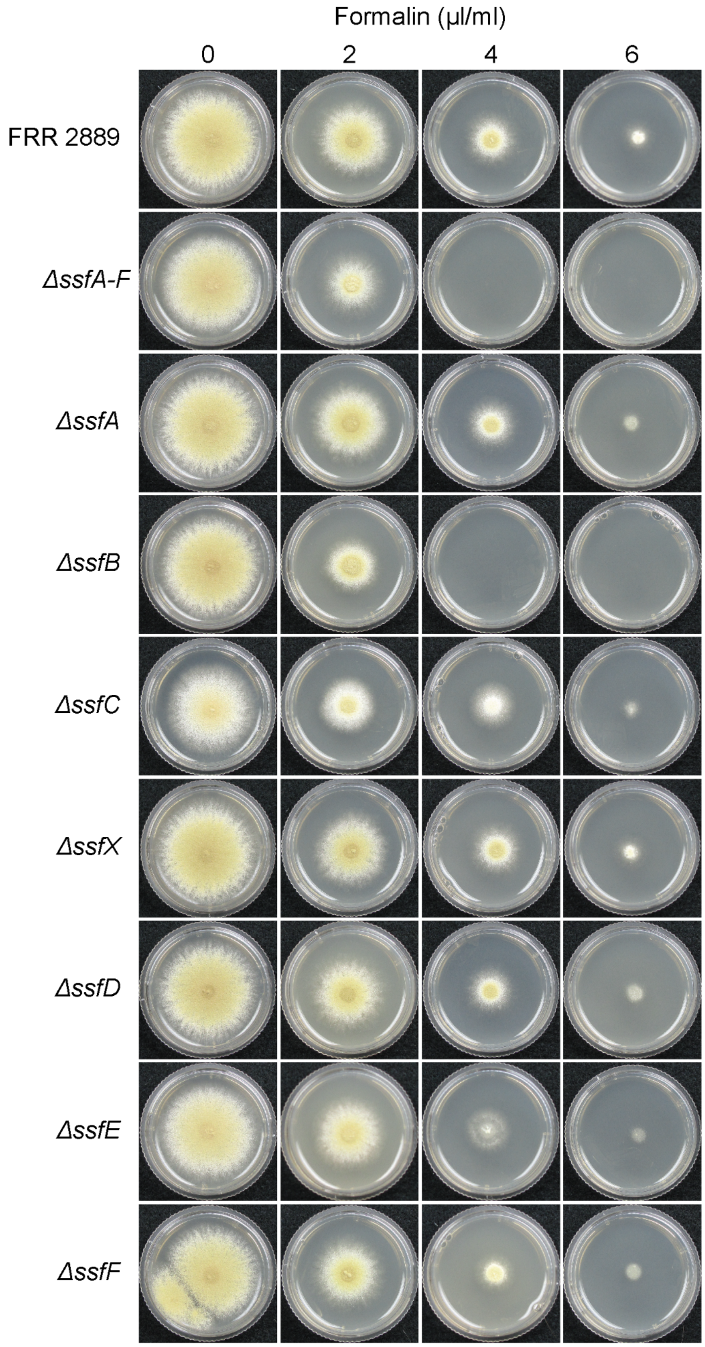


**Figure S8:** Wild type *P. variotii* FRR 2889 and 8 *ssf* gene mutant strains grown on varying concentrations of formaldehyde for 3 days at 30°C.

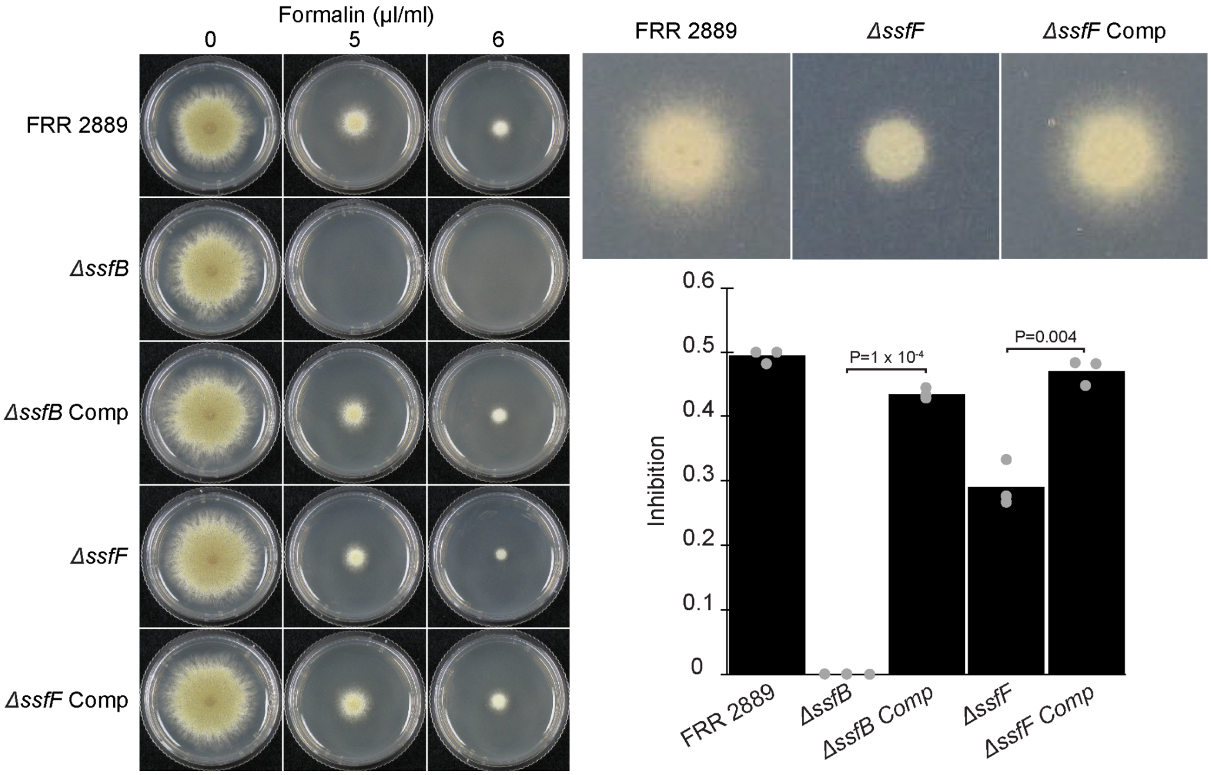


**Figure S9:** Growth rate of the *ΔssfB* and *ΔssfF* mutants compared to wild type and the corresponding genetically complemented counterparts. Plates grown for 3 days at 30°C.

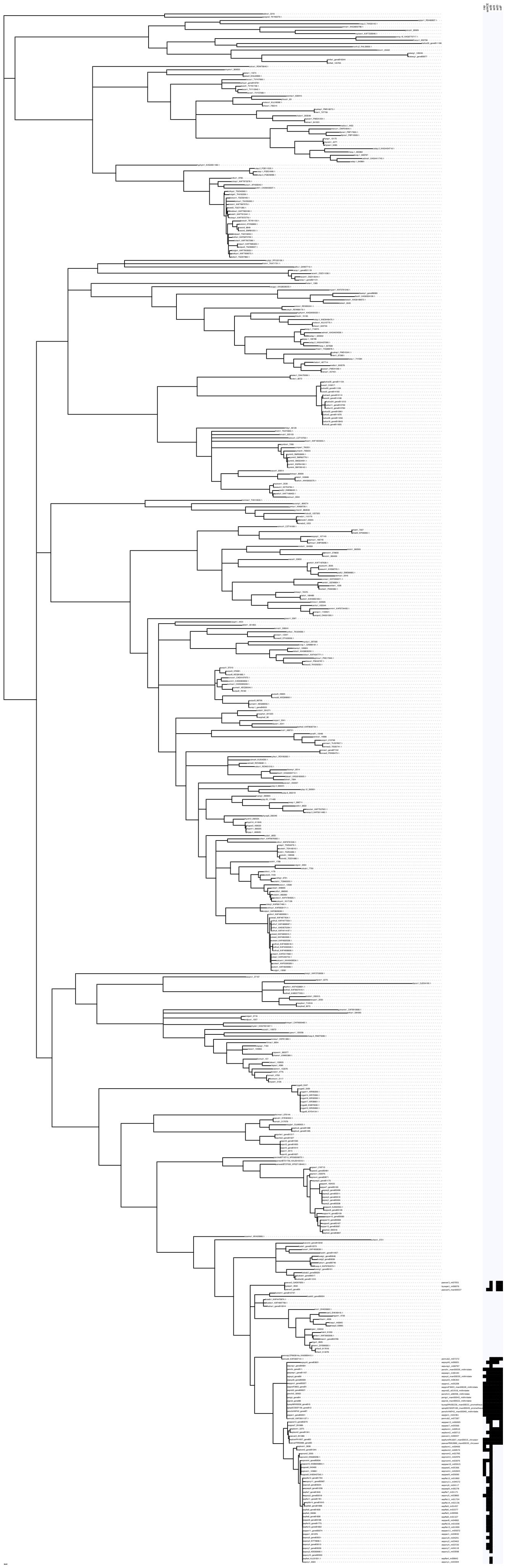

**Figure S10:** A midpoint-rooted maximum likelihood phylogeny of 468 SsfB sequences retrieved from the 2,899 genome database. Branch support was assessed with 1000 SH-aLRT tests and 1000 UFboot replicates. Branches with SH-aLRT support < 80% and UFboot support < 95% have been collapsed. *Starship* and gene neighborhood identifier codes are displayed to the right of sequences found in that associated region, for all sequences found to be a part of either a gene neighborhood or *Starship*. Neighborhoods were defined as containing homologs to at least 2 ssf genes of interest and Starships were either manually annotated or retrieved from the *Starship* database (Methods). To the right of the tree is a heatmap displaying the presence/absence of homologs to *Starship*- and ssf-associated genes in the *Starships* and gene neighborhoods associated with each sequence.

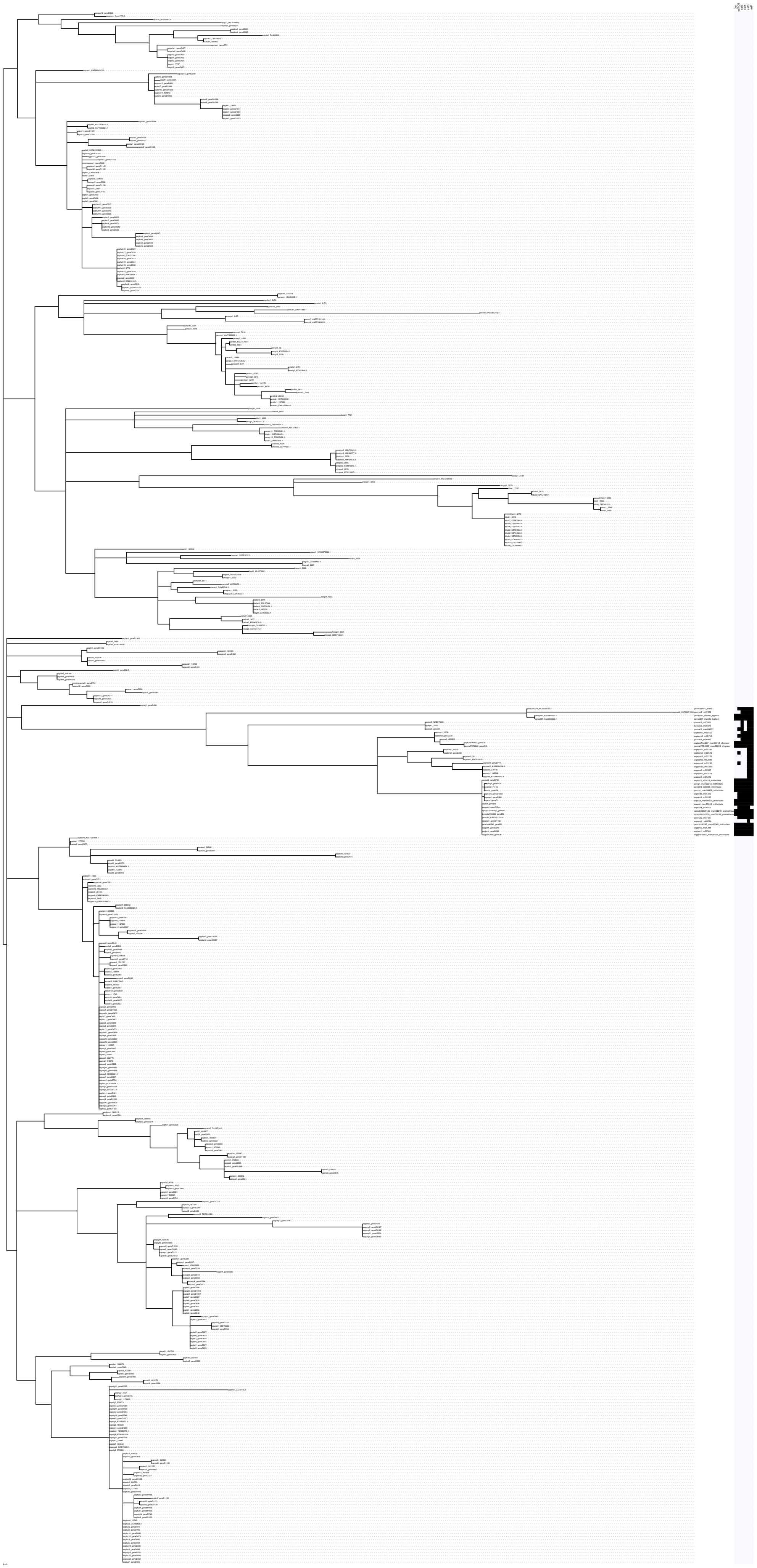

**Figure S11:** A midpoint-rooted maximum likelihood phylogeny of 484 SsfD sequences retrieved from the 2,899 genome database. Branch support was assessed with 1000 SH-aLRT tests and 1000 UFboot replicates. Branches with SH-aLRT support < 80% and UFboot support < 95% have been collapsed. *Starship* and gene neighborhood identifier codes are displayed to the right of sequences found in that associated region, for all sequences found to be a part of either a gene neighborhood or *Starship*. Neighborhoods were defined as containing homologs to at least 2 *ssf* genes of interest and Starships were either manually annotated or retrieved from the Starship database (Methods). To the right of the tree is a heatmap displaying the presence/absence of homologs to *Starship*- and *ssf*-associated genes in the *Starships* and gene neighborhoods associated with each sequence.

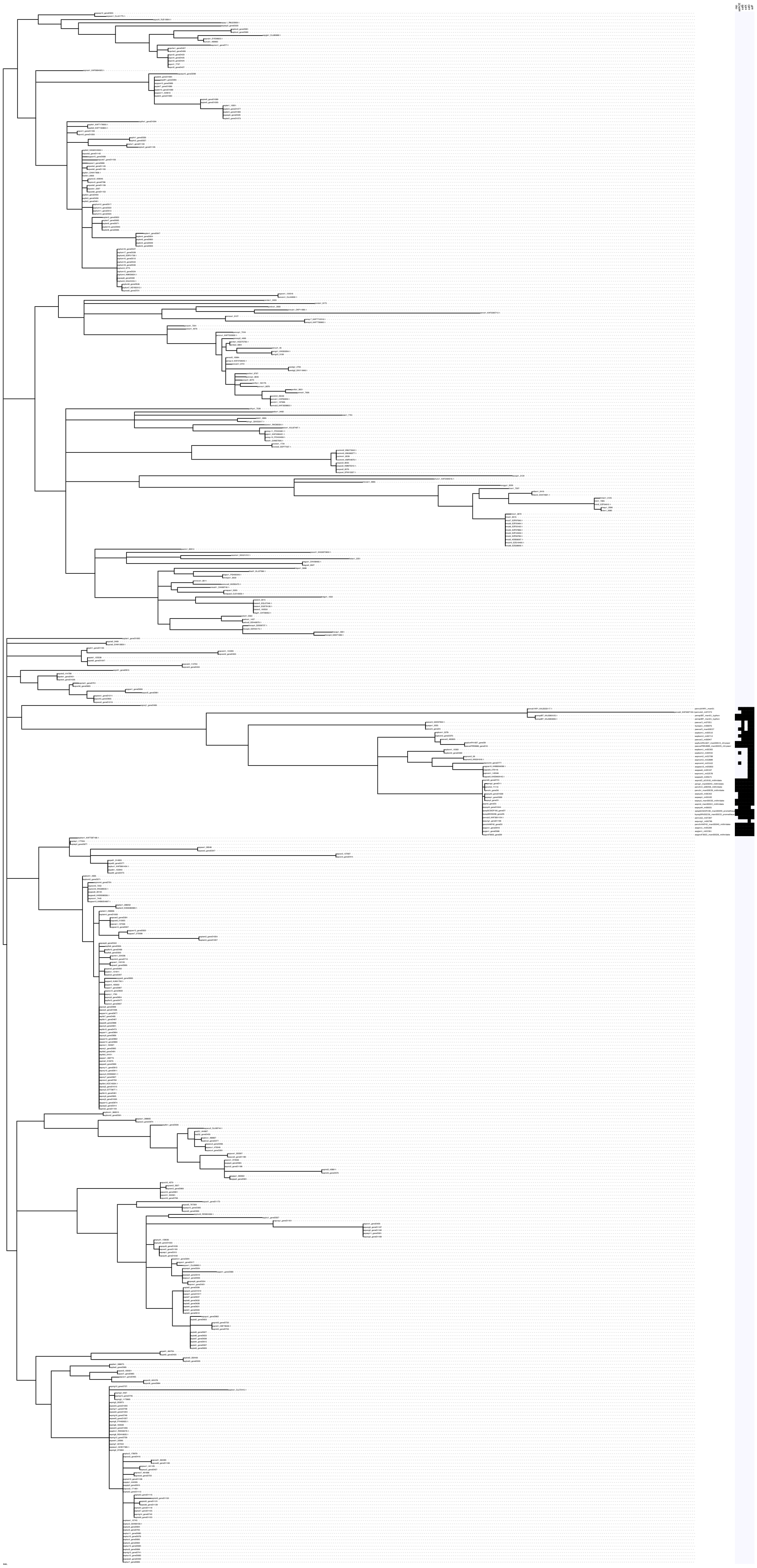

**Figure S12:** A midpoint-rooted maximum likelihood phylogeny of 124 SsfF sequences retrieved from the 2,899 genome database. Branch support was assessed with 1000 SH-aLRT tests and 1000 UFboot replicates. Branches with SH-aLRT support < 80% and UFboot support < 95% have been collapsed. *Starship* and gene neighborhood identifier codes are displayed to the right of sequences found in that associated region, for all sequences found to be a part of either a gene neighborhood or *Starship*. Neighborhoods were defined as containing homologs to at least 2 ssf genes of interest and *Starships* were either manually annotated or retrieved from the *Starship* database (Methods). To the right of the tree is a heatmap displaying the presence/absence of homologs to *Starship*- and *ssf*-associated genes in the *Starships* and gene neighborhoods associated with each sequence.

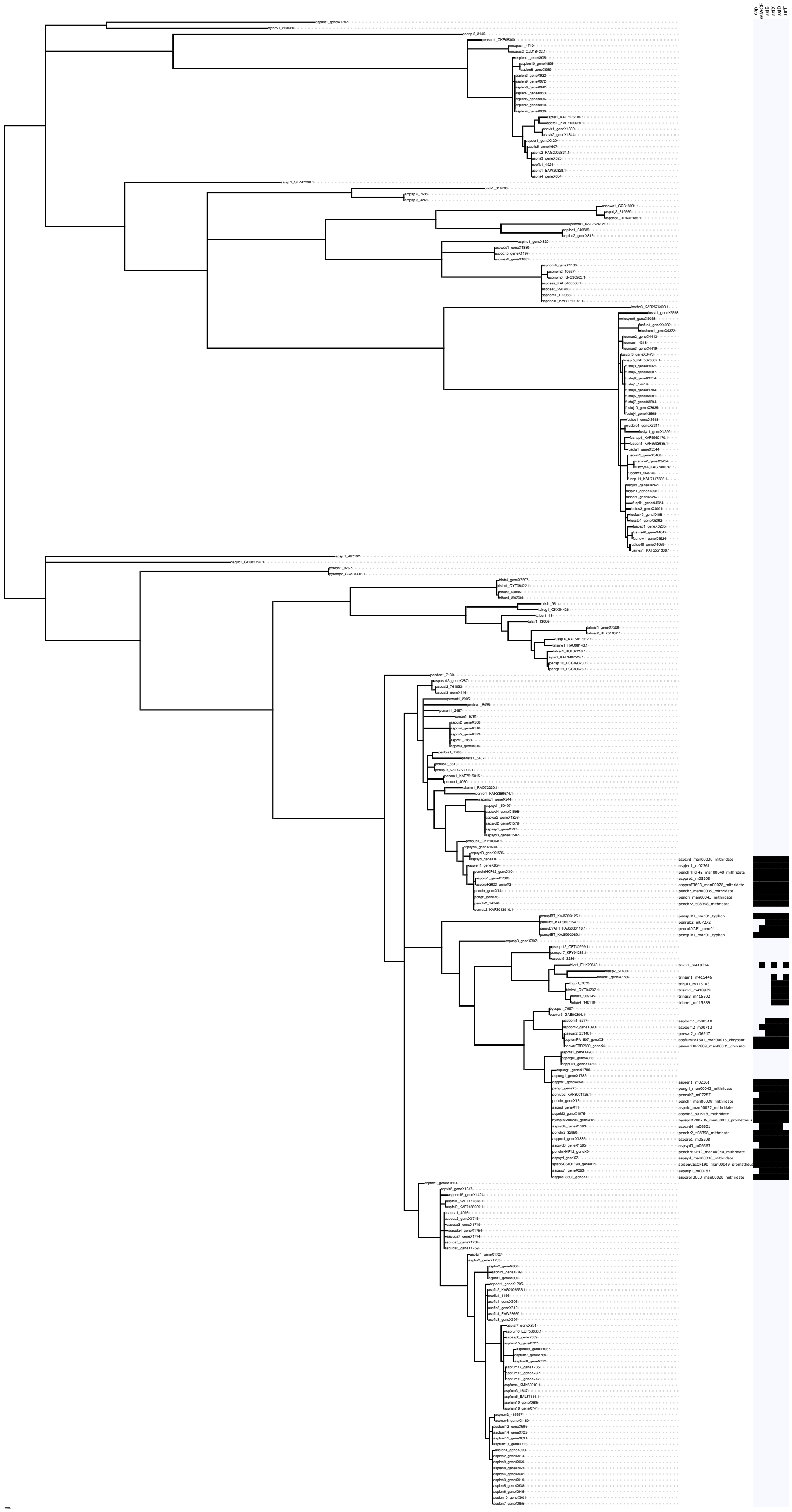
cap
ssfACE
ssfB
ssfX
ssfD
ssfF

**Figure S13:** A midpoint-rooted maximum likelihood phylogeny of 248 SsfX sequences retrieved from the 2,899 genome database. Branch support was assessed with 1000 SH-aLRT tests and 1000 UFboot replicates. Branches with SH-aLRT support < 80% and UFboot support < 95% have been collapsed. *Starship* and gene neighborhood identifier codes are displayed to the right of sequences found in that associated region, for all sequences found to be a part of either a gene neighborhood or *Starship*. Neighborhoods were defined as containing homologs to at least 2 ssf genes of interest and *Starships* were either manually annotated or retrieved from the *Starship* database (Methods). To the right of the tree is a heatmap displaying the presence/absence of homologs to *Starship*- and *ssf*-associated genes in the *Starships* and gene neighborhoods associated with each sequence.

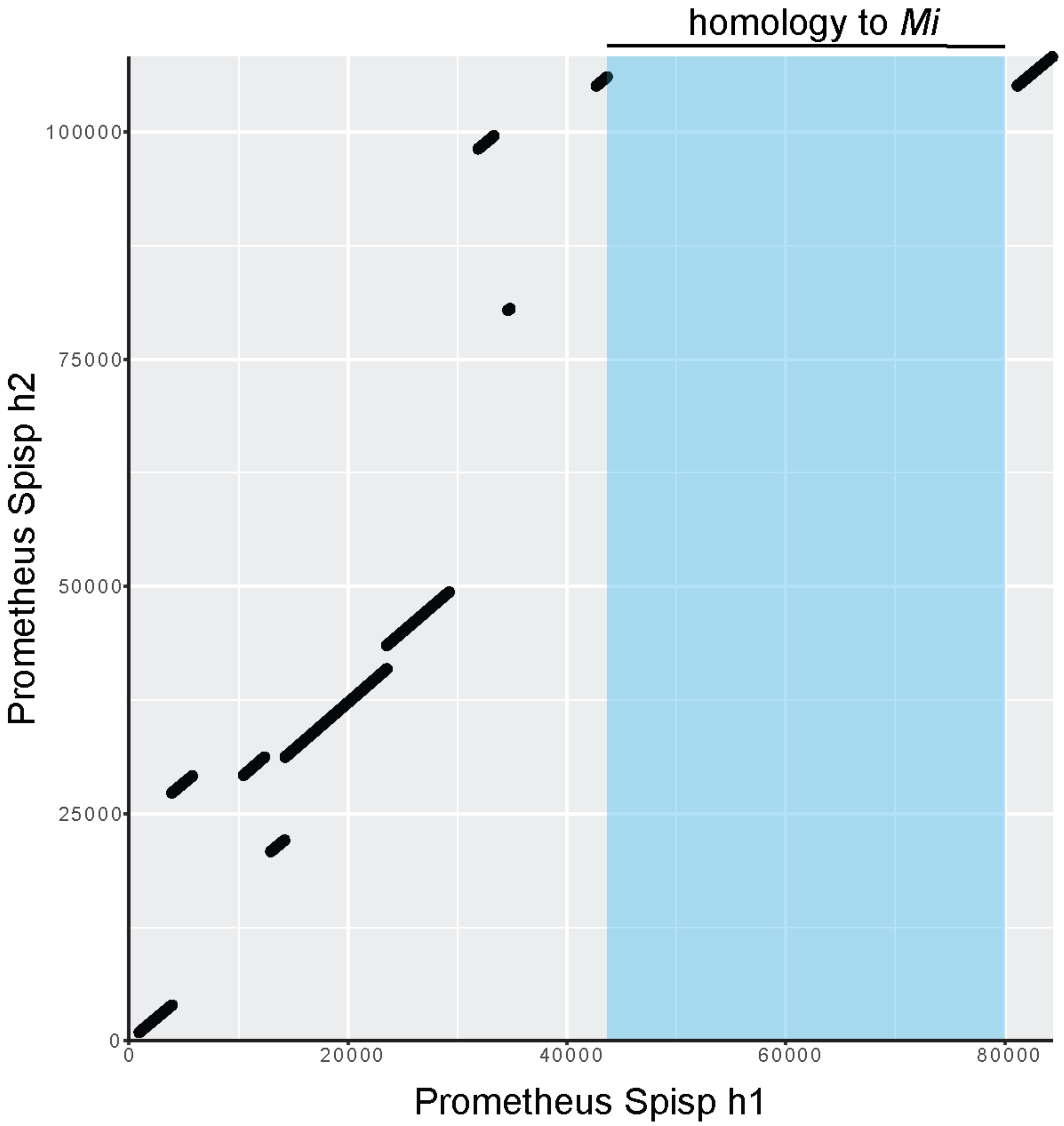


**Figure S14:** Dot plot comparing the two *Πρ* haplotypes in *Spiromastix sp*. SCSIOF190. Only haplotype 1 contains a large region of homology to *Mi* including the *ssf* cluster, this region is shaded blue.

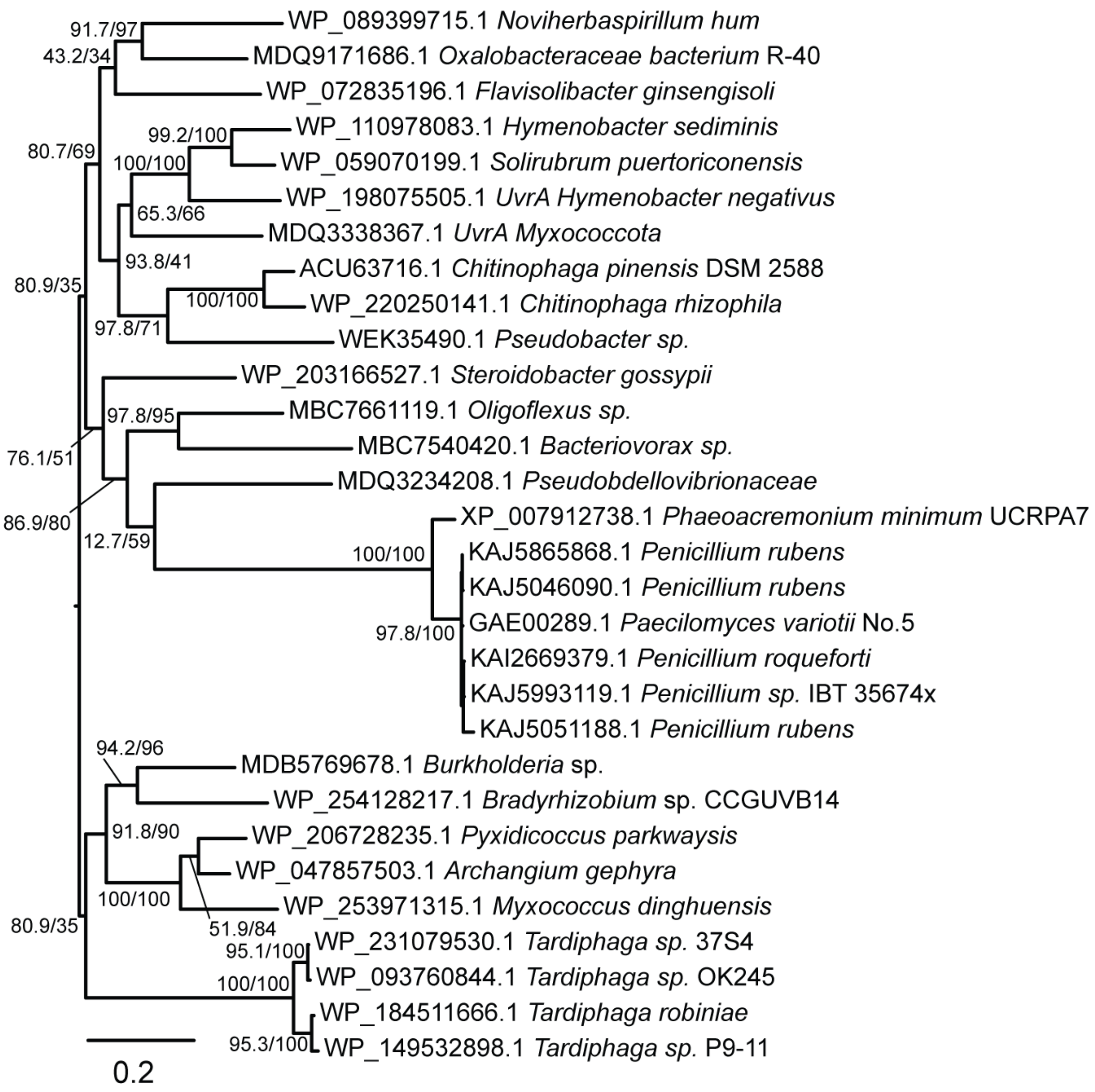


**Figure S15:** Protein phylogeny of a selection of closest homologs to the UvrA homolog encoded in certain ssf cluster-containing *Starships*. Sequences were aligned in MAFFT and tree generated using the default settings of IQ-TREE.

### Supplementary Tables

**Table S3:**

| Construct | Amplicon | Primer 1 | Primer 2 | Template |
|---|---|---|---|---|
| *ssfA-F* GFP replacement | 5´ flank of *ssfF* | AUB656 | AUB657 | FRR 2889 gDNA |
| | GFP-term-HYGR | AUB658 | AUB608 | PLAUB69 [29] |
| | 3´ flank of *ssfA* | AUB609 | AUB610 | FRR 2889 gDNA |
| *ssfA* GFP replacement | 5´ flank of gene | AUB605 | AUB606 | FRR 2889 gDNA |
| | GFP-term-HYGR | AUB607 | AUB608 | PLAUB69 [29] |
| | 3´ flank of gene | AUB609 | AUB610 | FRR 2889 gDNA |
| *ssfB* GFP replacement | 5´ flank of gene | AUB611 | AUB612 | FRR 2889 gDNA |
| | GFP-term-HYGR | AUB613 | AUB614 | PLAUB69 [29] |
| | 3´ flank of gene | AUB615 | AUB616 | FRR 2889 gDNA |
| *ssfC* GFP replacement | 5´ flank of gene | AUB617 | AUB618 | FRR 2889 gDNA |
| | GFP-term-HYGR | AUB619 | AUB620 | PLAUB69 [29] |
| | 3´ flank of gene | AUB621 | AUB622 | FRR 2889 gDNA |
| *ssfD* GFP replacement | 5´ flank of gene | AUB623 | AUB624 | FRR 2889 gDNA |
| | GFP-term-HYGR | AUB625 | AUB626 | PLAUB69 [29] |
| | 3´ flank of gene | AUB627 | AUB628 | FRR 2889 gDNA |
| *ssfE* GFP replacement | 5´ flank of gene | AUB629 | AUB630 | FRR 2889 gDNA |
| | GFP-term-HYGR | AUB631 | AUB632 | PLAUB69 [29] |
| | 3´ flank of gene | AUB633 | AUB634 | FRR 2889 gDNA |
| *ssfF* GFP replacement | 5´ flank of gene | AUB656 | AUB657 | FRR 2889 gDNA |
| | GFP-term-HYGR | AUB658 | AUB659 | PLAUB69 [29] |
| | 3´ flank of gene | AUB660 | AUB661 | FRR 2889 gDNA |
| *ssfX* GFP replacement | 5´ flank of gene | AUB727 | AUB728 | FRR 2889 gDNA |
| | GFP-term-HYGR | AUB729 | AUB730 | PLAUB69 [29] |
| | 3´ flank of gene | AUB731 | AUB732 | FRR 2889 gDNA |

| | | | | |
|---|---|---|---|---|
| *psfD* Knockout | 5´ flank of gene | AUB701 | AUB702 | FRR 2889 gDNA |
| | G418R | AUB703 | AUB704 | pMAI2 [96] |
| | 3´ flank of gene | AUB705 | AUB706 | FRR 2889 gDNA |
| *psfF* Knockout | 5´ flank of gene | AUB707 | AUB708 | FRR 2889 gDNA |
| | G418R | AUB709 | AUB710 | pMAI2 [96] |
| | 3´ flank of gene | AUB711 | AUB712 | FRR 2889 gDNA |
| *ssfF* comp | Wildtype copy of gene | AUB755 | AUB756 | FRR 2889 gDNA |
| *ssfB* comp | Wildtype copy of gene | AUB699 | AUB700 | FRR 2889 gDNA |

**Table S4:**

| **Transformant** | **Upstream primer** | **Downstream primer** |
|---|---|---|
| *ssfA* GFP replacement | AUB666 | AUB667 |
| *ssfB* GFP replacement | AUB670 | AUB671 |
| *ssfC* GFP replacement | AUB674 | AUB675 |
| *ssfD* GFP replacement | AUB678 | AUB679 |
| *ssfE* GFP replacement | AUB682 | AUB683 |
| *ssfF* GFP replacement | AUB686 | AUB687 |
| *psfD* Knockout | AUB719 | AUB720 |
| *psfF* Knockout | AUB723 | AUB724 |
| *ssfX* GFP replacement | AUB741 | AUB681 |
